# ProClust provides reliable statistical significance of clusters for single-cell transcriptomic data

**DOI:** 10.64898/2026.11.08.764985

**Authors:** Haina Qian, Haoze Wu, Yifan Dai, Guanao Yan, Yufeng Liu

## Abstract

Clustering of single-cell RNA sequencing (scRNA-seq) data is widely used to identify cell populations. However, overclustering can partition a single cell population into multiple clusters, potentially leading to spurious identification of cell subpopulations. Assessing the statistical significance of clusters is therefore needed to support their biological interpretation. We propose ProClust, a statistical test for clustering significance in scRNA-seq data. ProClust defines a single cluster as a log-concave distribution and adopts a projection-based approach to quantify clustering uncertainty through *p*-values. These projections can also provide interpretable visual evidence for individual clusterings. ProClust merges clusters when there is insufficient evidence to reject the single-cluster null hypothesis, refining an initial overclustering without prespecifying the final number of clusters. Across parametric and PBMC-based semi-synthetic simulations, ProClust avoided false subdivisions under single-population null settings while consistently recovering the underlying population structure across different numbers of true populations. In real scRNA-seq datasets from the Human Lung Cell Atlas, Tabula Muris, and *Hydra*, ProClust produced clustering solutions that showed overall agreement with reference annotations while retaining supported substructure and merging groups with insufficient evidence for separation. Together, these results demonstrate that ProClust provides a statistically principled and interpretable framework for refining an initial overclustering and distinguishing supported cellular heterogeneity. The ProClust R package is available at https://github.com/Anonym027/ProClust.

## 1 Introduction

Single-cell RNA sequencing (scRNA-seq) enables biological heterogeneity to be characterized at the resolution of individual cells and has become an important tool in modern genomics (Nesari et al., 2026; Jovic et al., 2022). These cellular-resolution measurements have facilitated the identification of previously unrecognized cell types and transient cellular states and supported increasingly detailed studies of development, tissue organization, and disease (Woo and Eyun, 2025). A central task in analyzing such data is to group cells with similar transcriptional profiles into clusters representing candidate biological populations (Stegle et al., 2015). These empirical groupings provide the basis for cell-type annotation, marker-gene identification, trajectory inference, differential expression analysis, and subsequent biological interpretation (Haque et al., 2017). The reliability of the inferred clusters is therefore important for downstream analyses: underclustering may obscure subtle but functionally relevant heterogeneity by combining distinct populations, whereas overclustering may produce divisions associated with technical noise or sampling variation that could be incorrectly interpreted as distinct cell types or states (Zappia and Oshlack, 2018; Petegrosso et al., 2020).

The cluster labels inferred from scRNA-seq data are data-dependent random objects, and their uncertainty should therefore be quantified. In practice, clustering workflows such as SC3 (Kiselev et al., 2017), Seurat (Satija et al., 2015; Stuart et al., 2019), and Monocle (Trapnell et al., 2014; Cao et al., 2019) can estimate candidate clustering results, while choices of clustering resolution or the number of clusters are commonly used to control the granularity of the resulting cellular populations (Xiang et al., 2021; Petegrosso et al., 2020). However, selecting a preferred resolution does not quantify the statistical evidence supporting the partition. It does not answer whether two inferred groups are sufficiently distinct to support their interpretation as separate populations, or whether the observed separation could plausibly arise from sampling variation within a single population. Indeed, clustering algorithms can produce seemingly well-separated partitions even for data generated from a single homogeneous distribution, such as a Gaussian distribution (Liu et al., 2008).

Statistical significance analysis for clustering has been developed under several specific single-cluster models, but important challenges remain unresolved for scRNA-seq data. The central difficulty is that any significance test must specify what constitutes a single cluster under the null hypothesis. Existing methods have largely relied on restrictive parametric assumptions (Liu et al., 2008; Huang et al., 2015; Kimes et al., 2017; Y. T. Chen and Witten, 2023; Gao et al., 2024), and only more recently have such ideas been adapted to single-cell applications (Grabski et al., 2023). For example, SigClust defines a single cluster as a Gaussian distribution (Liu et al., 2008), with extensions to count data (Dai et al., 2025), while scSHC adopts a Poisson log-normal model for scRNA-seq data (Kimes et al., 2017; Grabski et al., 2023). This is particularly limiting for scRNA-seq data, whose observed distributions can vary substantially across sequencing platforms, sequencing depths, and biological settings (Tian et al., 2019; W. Chen et al., 2021). More importantly, these assumptions are difficult to assess in practice. Once a candidate cluster has been observed, it is generally difficult to determine whether deviations from Gaussian or Poisson-like behavior indicate genuine clustering structure or simply mis-specification of the assumed model. This makes a small *p*-value from these methods difficult to interpret. CHOIR instead evaluates candidate separations using random-forest classification and permutation testing (Sant et al., 2025), but does not provide distributional characterization of a homogeneous cluster. As we show later, this lack of a flexible and interpretable single-cluster model can lead existing methods to overcluster and produce false discoveries under modest departures from their assumed null distributions.

To address these challenges, we propose ProClust, a flexible framework for assessing the statistical significance of clusters in single-cell data. ProClust characterizes a single homogeneous population using the family of log-concave distributions, providing a broad unimodal null model that accommodates skewness and dependence. Log-concave distributions have long been considered as a natural model for a single cluster (Bagnoli and Bergstrom, 2005; Samworth, 2018; Dai et al., 2026) and include many distributions commonly used for scRNA-seq data, such as negative binomial and multinomial distributions (Townes et al., 2019; Sun et al., 2021). However, directly estimating a high-dimensional log-concave distribution remains challenging (Cule et al., 2010; Samworth, 2018). Motivated by the recent hunt-and-test procedure for unimodality testing (Guo and Shah, 2025), ProClust instead first finds a low-dimensional clustering subspace that captures the proposed clustering structure, and then performs statistical inference within this subspace. This approach avoids costly and potentially inaccurate estimation of log-concave densities while retaining the clustering structure.

ProClust advances statistical inference for single-cell clustering in three respects. First, ProClust is model-free and robust to model misspecification, characterizing cluster homogeneity through the broad class of log-concave distributions. This formulation provides a substantially broader family of single-cluster models than conventional Gaussian and Poisson assumptions, and thereby enables significance assessment across diverse single-cell data. Second, ProClust identifies the low-dimensional structure underlying clustering significance. As shown later, *K* candidate clusters can be represented in a *K* − 1-dimensional subspace without losing the clustering structure relevant to the test. This characterization complements formal hypothesis testing with interpretable low-dimensional visualization of candidate splits. These projections reveal the underlying clustering structure, allowing the evidence for retaining or merging candidate groups to be examined visually. Third, across the settings considered in this study, ProClust yields well-calibrated *p*-values, maintains power for detecting subtle population separations, accurately recovers underlying clustering structure, and achieves substantial computational gains relative to resampling-intensive significance-based procedures. Its computational design supports application to increasingly large single-cell datasets. In the following sections, we first evaluate the statistical calibration, power, clustering performance, and runtime of ProClust across comprehensive simulations, and then examine its clustering results and biological interpretability across three scRNA-seq atlases.

### 2 Results

### 2.1 Overview of ProClust Algorithm

ProClust is a projection-based hierarchical clustering framework that combines discriminative dimension reduction with statistical hypothesis testing. Starting from an initial clustering or a set of reference annotations, ProClust constructs a hierarchical cluster tree and recursively evaluates candidate splits in a top-down manner. At each internal node, ProClust tests whether the proposed partition is supported by sufficient statistical evidence and assigns a corresponding *p*-value. Supported partitions are retained, and their descendant nodes are evaluated recursively, whereas unsupported splits are merged. In this way, ProClust determines the final clustering based on statistically supported partitions without requiring the number of clusters to be specified in advance. An overview of the ProClust workflow is shown in Fig. 1.

**Figure 1:**
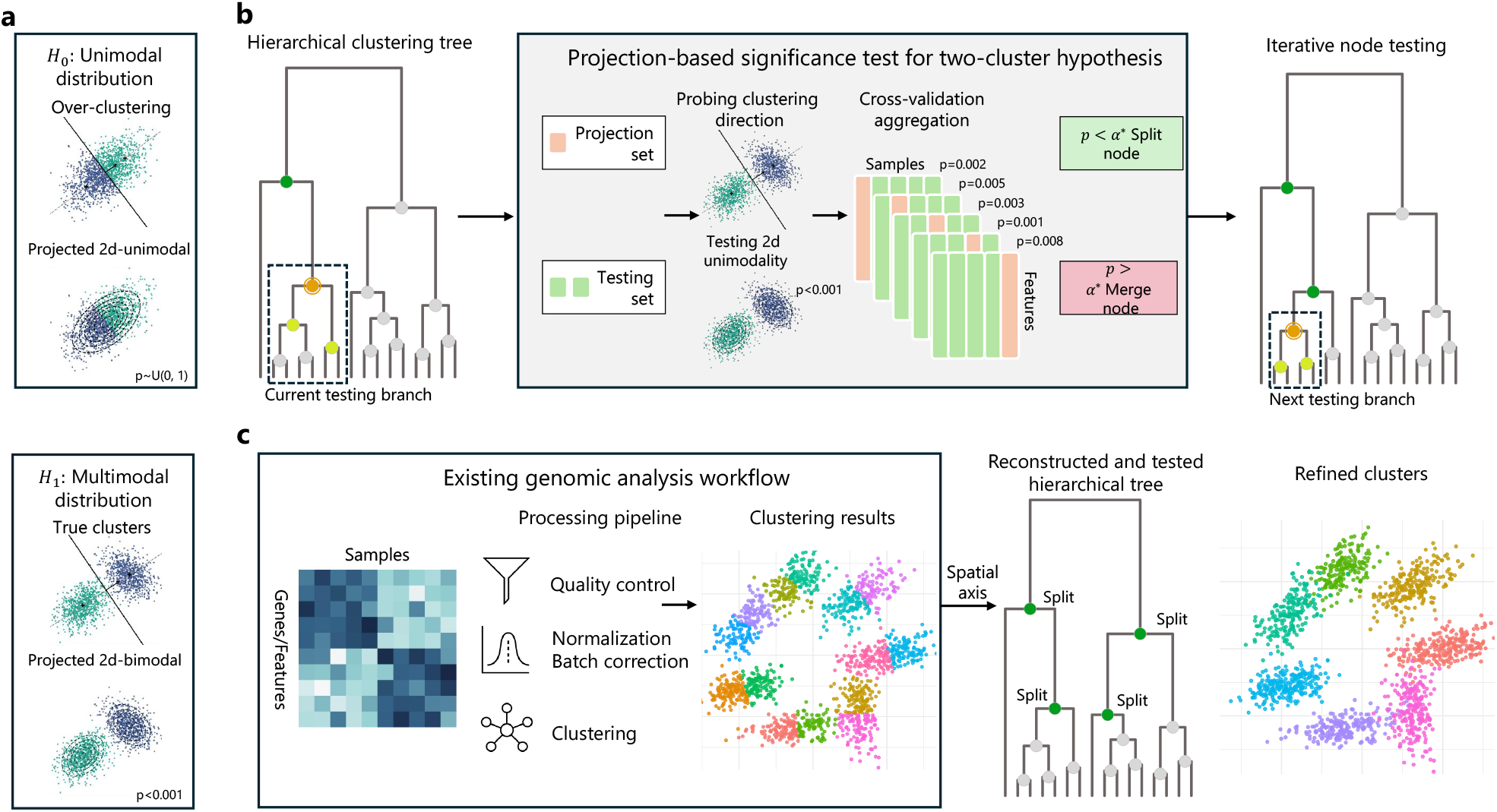
Workflow of ProClust: a significance testing framework for genomic clustering. **a**. *Modality-based definition of clusters*. Under the null hypothesis of a single cluster, the projection of data onto a two-dimensional subspace yields a unimodal distribution; under the alternative, the projected distribution becomes at least bimodal. **b**. *Testing procedure of ProClust*. ProClust begins with a hierarchical clustering tree and iteratively evaluates the statistical significance of each split from top to bottom. For each node, the data are divided into a projection set to estimate the 2D clustering direction and a testing set to assess 2D unimodality. The resulting *p*-values are integrated via cross-validation and aggregated across random restarts via the median. Tree-level family-wise error rate is controlled using a node-specific threshold *α*^*\**^, with nodes split or merged based on whether the median *p*-value falls below or exceeds *α*^*\**^. **c**. *Integration with genomic analysis pipelines*. Given a processed genomic count matrix, ProClust can be seamlessly applied to assign statistical *p*-values to clusters by reconstructing the hierarchical tree and iteratively refining over-clustered populations.

To assess the significance of candidate cluster divisions, ProClust exploits two key properties of clustering structure. First, clustering contrasts are intrinsically low-dimensional. For *K* clusters, the between-cluster structure can be represented in a subspace of dimension at most *K* − 1 (Fig. 2a). Because ProClust evaluates local splits within a hierarchical tree, only a small number of candidate clusters, typically *K* = 2 or 3, need to be considered at each node, yielding a one- or two-dimensional discriminative subspace. ProClust uses local Fisher discriminant analysis (LFDA; Sugiyama, 2007), a supervised linear dimension-reduction method, to estimate this subspace while preserving local within-cluster structure. Second, ProClust avoids restrictive parametric assumptions on the distribution of a homogeneous cluster. Rather than assuming Gaussian, Poisson, or another prespecified parametric family, ProClust models a single cluster using a log-concave distribution, providing a flexible class of unimodal distributions (Fig. 2b). The observed cluster separation is then compared with the separation obtained from simulated data under the fitted log-concave null distribution. A candidate split is considered significant only when its observed separation is sufficiently stronger than would be expected under this single-cluster null (Fig. 2c).

**Figure 2:**
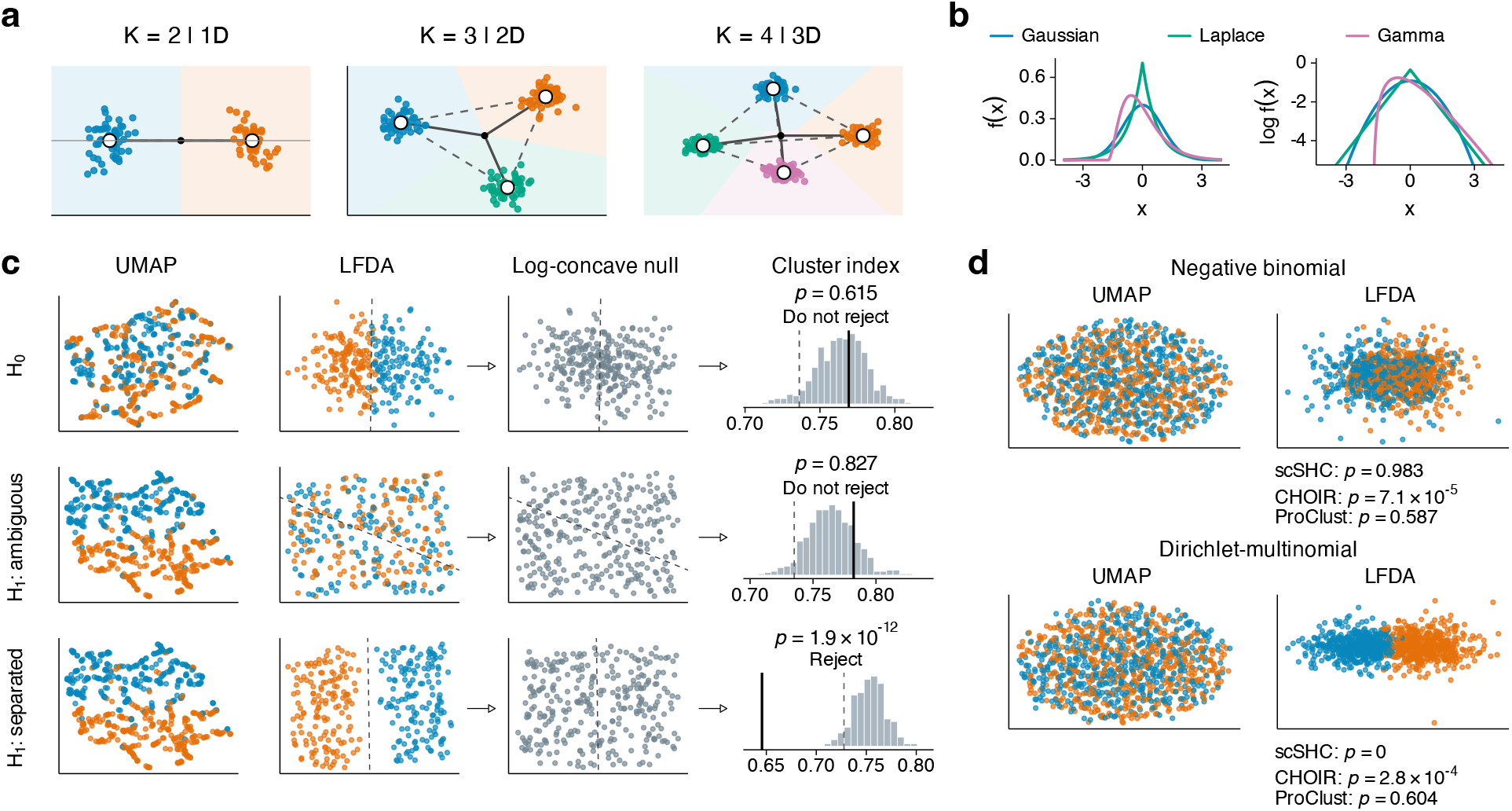
Key concepts underlying ProClust. a. Between-cluster contrasts are low-dimensional. For *K* clusters, the centered cluster centroids span a subspace of dimension at most *K* − 1, allowing significance testing to focus on the corresponding discriminative structure. **b**. The log-concave family includes diverse unimodal distributions, including Gaussian, Laplace, and Gamma distributions with shape parameter at least one. **c**. ProClust tests whether the observed cluster index is significantly smaller than expected under a fitted null distribution with log-concave margins. The three rows illustrate a single cluster (*H*_0_), multiple clusters with incorrect fitted labels (*H*_1_ : ambiguous), and multiple clusters with correct fitted labels (*H*_1_ : separated). The latter two share the same underlying distribution but differ in fitted labels. Only the last case leads to rejection. **d**. UMAP and LFDA embeddings of negative binomial and Dirichlet–multinomial simulations generated without group-wise mean differences.

ProClust *p*-values are accompanied by LFDA embeddings that allow researchers to visually examine the clustering structure underlying each hypothesis test (Fig. 2c). Because LFDA uses only linear projections, its geometry remains directly interpretable and is less prone to introducing artificial separation than nonlinear visualization methods such as UMAP (Becht et al., 2019). At the same time, ProClust embeddings can recover the between-cluster structure of genuinely distinct groups, including weakly separated clusters that may be obscured in UMAP visualizations. Notably, LFDA in ProClust is designed to facilitate examination of between-cluster contrasts involving a small number of groups, typically two or three clusters. It is not intended to replace UMAP for visualization of a complete single-cell atlas containing many cell populations.

The nonparametric formulation makes ProClust less sensitive to distributional misspecification, which can otherwise lead existing parametric testing procedures to spuriously reject cluster homogeneity. For example, negative binomial and Dirichlet–multinomial distributions with 1,000 features produce similar UMAP and LFDA embeddings after a standard scRNA-seq preprocessing pipeline. Both embeddings show no apparent clustering structure. However, CHOIR (Sant et al., 2025) rejects cluster homogeneity under both distributions, while scSHC (Grabski et al., 2023) rejects homogeneity under the Dirichlet–multinomial distribution. In contrast, ProClust does not reject cluster homogeneity under either setting.

### 2.2 ProClust reliably recovers population structure across simulation frameworks

We evaluated ProClust using two complementary simulation frameworks. The parametric simulations provided a controlled setting in which the number of ground-truth populations and the sources of within-population variation were explicitly specified, whereas the PBMC-based semi-synthetic simulations preserved population-specific expression characteristics and gene–gene dependence estimated from real single-cell data. Despite these distinct data-generating mechanisms, the two frameworks yielded broadly consistent performance patterns under both the null and alternative hypotheses (Fig. 3a–c).

**Figure 3:**
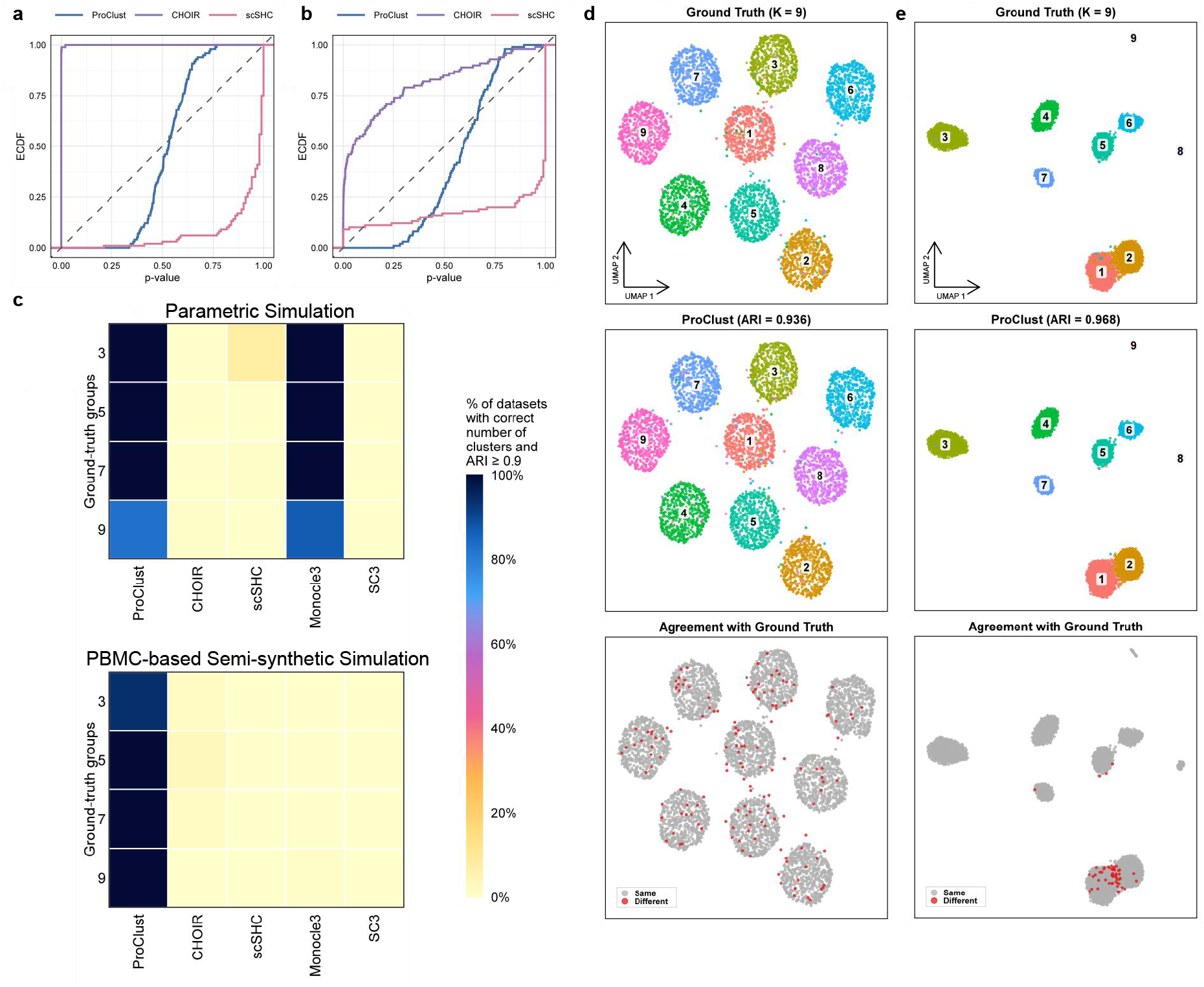
Evaluation of statistical calibration and clustering recovery across simulation frameworks. **a–b**, Empirical cumulative distribution functions (ECDFs) of *p*-values under the single-population null hypothesis (*H*_0_) for ProClust, CHOIR, and scSHC in the parametric (**a**) and PBMC-based semi-synthetic (**b**) simulations. The dashed diagonal indicates the theoretical cumulative distribution function of the Uniform(0, 1) distribution. **c**, Joint recovery performance under the alternative hypothesis (*H*_1_) across *K*_true_ = 3, 5, 7, and 9. Colors indicate the percentage of simulation replicates in which each method simultaneously recovered the correct number of populations and achieved an adjusted Rand index (ARI) of at least 0.9. Results are shown separately for the parametric (top) and PBMC-based semi-synthetic (bottom) simulation frameworks. **d–e**, Representative datasets with *K*_true_ = 9 from the parametric (**d**) and PBMC-based semi-synthetic (**e**) simulations. From top to bottom, panels show the ground-truth populations, ProClust clustering results with the corresponding ARI, and cell-level agreement between the ProClust assignments and ground-truth labels. Gray points indicate cells assigned consistently with the ground truth, whereas red points indicate disagreements.

Under the null hypothesis, ProClust produced no *p*-values below the prespecified significance threshold of 0.05 in either simulation framework, corresponding to an observed type I error rate of 0%, despite S-shaped empirical *p*-value distributions that deviated from uniformity. In contrast, the CHOIR *p*-values showed a pronounced concentration near zero, resulting in substantial inflation of the type I error rate. Although the scSHC *p*-values were predominantly concentrated near one in both simulation frameworks, scSHC was conservative at the 0.05 significance level in the parametric simulation but anticonservative in the PBMC-based semi-synthetic simulation.

Under the alternative hypothesis, we evaluated whether each method simultaneously recovered the ground-truth number of populations and produced cluster assignments closely aligned with the ground-truth labels. Successful recovery was defined as identifying exactly *K*_true_ clusters while attaining an adjusted Rand index (ARI) of at least 0.9 (Fig. 3c). ProClust achieved consistently high joint recovery rates across *K*_true_ = 3, 5, 7, and 9 in both simulation frameworks.

In the parametric simulations, ProClust recovered the exact number of populations in at least 97% of replicates at every value of *K*_true_. Monocle3 also achieved high ARI values but was less reliable in recovering the ground-truth number of populations overall, particularly at *K*_true_ = 9, where its exact recovery rate was 87%, compared with 97% for ProClust. CHOIR, scSHC, and SC3 rarely satisfied both recovery criteria simultaneously.

The advantage of ProClust was more pronounced in the PBMC-based semi-synthetic simulations, where it achieved the highest joint recovery rate at every value of *K*_true_. ProClust recovered the exact number of populations in 94% of replicates at *K*_true_ = 3 and in all replicates at *K*_true_ = 5, 7, and 9. By contrast, scSHC, SC3 and CHOIR overestimated the number of populations, whereas Monocle3 consistently underestimated it.

Representative datasets with *K*_true_ = 9 further illustrate these overall patterns (Fig. 3d–e). In both the parametric and PBMC-based semi-synthetic simulations, the clusters returned by ProClust closely reproduced the nine ground-truth populations. Direct comparison of the ProClust assignments with the ground-truth labels showed agreement for the majority of cells, with relatively few mismatched assignments. Together, these results indicate that ProClust can terminate further subdivision when no additional population structure is present while retaining the ability to recover multiple populations across distinct data-generating mechanisms.

### 2.3 ProClust recovers major immune cell populations and reveals alveolar macrophage heterogeneity in the HLCA

We evaluated ProClust on a subset of the Human Lung Cell Atlas (HLCA) core dataset, comprising immune cells from healthy human lung tissue with expert-curated level-4 subtype annotations (Sikkema et al., 2023).

We first generated an over-clustered initial partition using Leiden (Traag et al., 2019) with a resolution parameter of 1.0, which served as the common input for two post-clustering testing methods: ProClust and scSHC. CHOIR, by contrast, builds its own hierarchical over-clustering internally as part of its standard pipeline, and was therefore run independently rather than on the shared Leiden partition. Clustering performance was evaluated using the Adjusted Rand Index (ARI) and Normalized Mutual Information (NMI) against the ground-truth cell-type annotations (Fig. 4). ProClust achieved the highest ARI and NMI among the three methods, indicating the best agreement with the reference annotations. Visual inspection of the UMAP embeddings further shows that ProClust produces a balanced clustering solution by preserving biologically meaningful cell populations while avoiding unnecessary fragmentation. In contrast, CHOIR merges some biologically distinct populations into coarser clusters, while scSHC over-clusters, closely mirroring the initial Leiden partition. These results demonstrate that ProClust provides a better balance between resolving cellular heterogeneity and preserving biologically meaningful structure.

**Figure 4:**
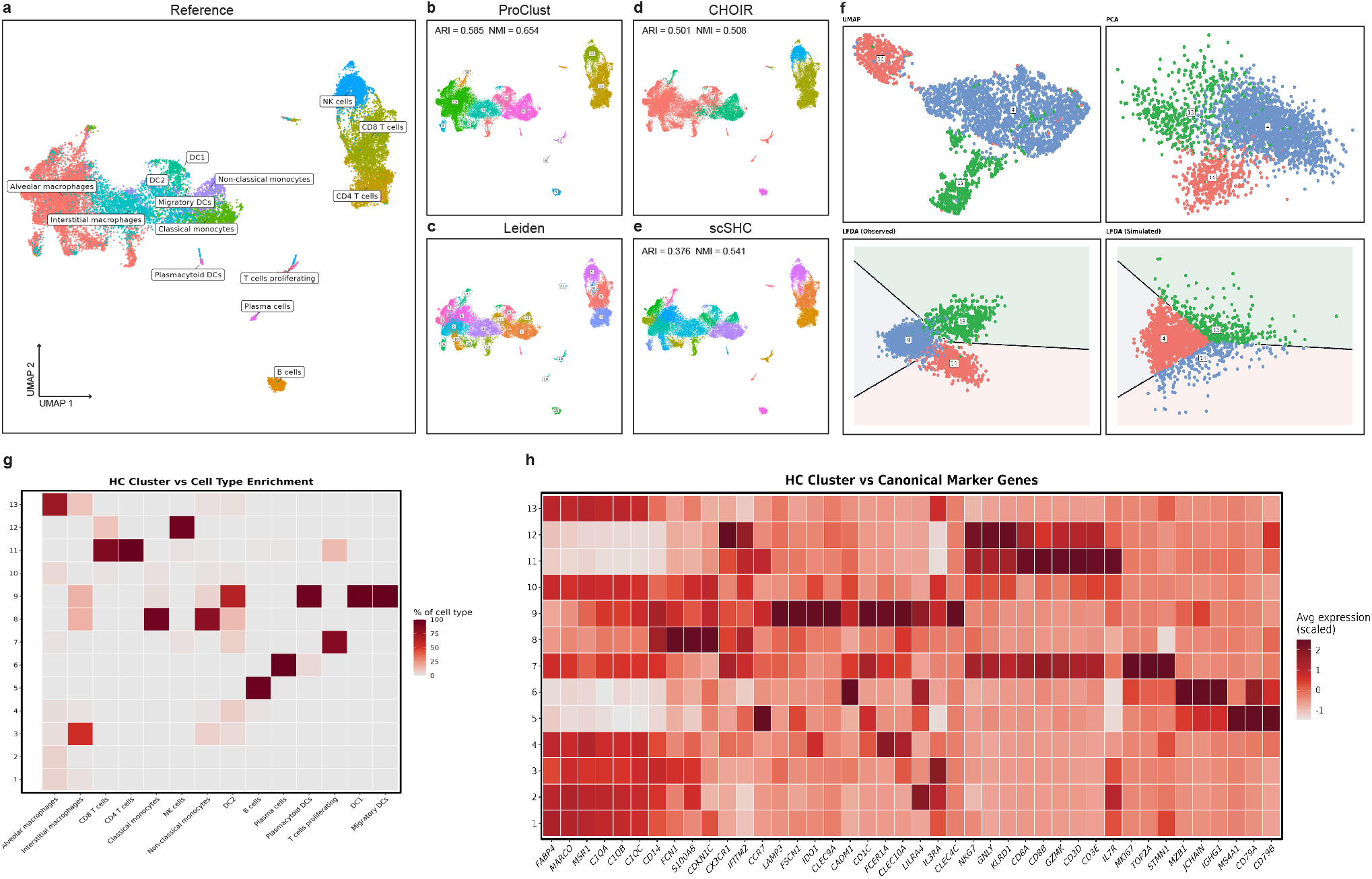
Comparison of clustering results and biological validation on the HLCA immune dataset. **a**, UMAP embedding of the sampled HLCA immune cells colored by the reference cell annotations. **b**, UMAP embeddings colored by the clusters identified by ProClust. **c**, The initial over-clustered partition (Leiden, resolution = 1.0) used as the common input for the testing procedures shown in **b** and **e. d**, UMAP embedding colored by the clusters identified by CHOIR. **e**, UMAP embedding colored by the clusters identified by scSHC. **f**, Visualization of the LFDA-based significance testing framework. The top row shows the UMAP and Harmony-PCA representations of the observed cells. The bottom row shows the LFDA projection of the observed data together with the decision regions (left) and simulated samples generated from the fitted null distribution within the same LFDA space (right). **g**, Heatmap showing, for each ground-truth cell type, the percentage of its cells that fall into each ProClust cluster. **h**, Heatmap showing the scaled average expression of canonical immune marker genes across the clusters identified by ProClust.

To further examine the statistical basis of these results, we visualized the significance testing procedure underlying ProClust using one representative split as an example (clusters 4, 13, and 14; Fig. 4f). The observed cells and simulated null samples were projected into the same LFDA space, enabling direct comparison between the observed cluster structure and the fitted null distribution: if the two distributions look similar, the split is not statistically supported and the clusters are merged; if the observed separation is sharper than what the null distribution produces, the split is retained as significant. Unlike conventional clustering methods, ProClust explicitly visualizes the decision regions used during significance testing, making the statistical decision process transparent and interpretable.

We next assessed the biological validity of the ProClust clusters using canonical marker genes (Fig. 4g, h). The clusters identified by ProClust were highly consistent with established immune cell identities: the B-cell cluster showed strong expression of MS4A1, CD79A, and CD79B (Hao et al., 2021), the plasma-cell cluster was characterized by high expression of MZB1, JCHAIN, and IGHG1 (Hao et al., 2021), the NK-cell cluster expressed the cytotoxic markers NKG7, GNLY, and KLRD1 (Hao et al., 2021), and the classical monocyte cluster displayed the expected markers CD14, FCN1, and S100A8 (Zilionis et al., 2019). Multiple clusters annotated as alveolar macrophages also consistently expressed the canonical alveolar macrophage markers FABP4, MARCO and MSR1 (Zilionis et al., 2019). Overall, these marker expression patterns agree well with the assigned cell identities, indicating that ProClust successfully recovered biologically meaningful immune cell populations.

Several clusters also showed enrichment for distinct marker modules, pointing to additional structure within the alveolar macrophage population beyond its broad annotation (see Supplementary Materials). Cluster 7 stood out with a clean proliferation signature (MKI67, TOP2A, STMN1, and CENPF) (Liégeois et al., 2022) that was largely absent from the other clusters. Cluster 2 combined strong inflammatory signal (GBP1 and IL1B) (Ma et al., 2022) with metallothionein expression (MT1E and MT2A) (Sauler et al., 2022), whereas cluster 10 showed a similar metallothionein-dominant profile but without the accompanying inflammatory signature. Cluster 1, meanwhile, was most enriched for homeostatic and lipid-associated markers (PLIN2 and MARCO) (Hurskainen et al., 2021). While some of these marker programs overlapped across clusters, the overall pattern suggests that the alveolar macrophage population comprises multiple substates rather than a single homogeneous group, and that finer subclustering could resolve additional biologically meaningful heterogeneity.

While ProClust accurately recovered most major immune cell populations, CD4 and CD8 T cells were not fully separated in the global clustering. Repeating the analysis on the full T-cell population using a targeted panel of T-cell lineage marker genes, rather than the genome-wide highly variable gene set, yielded a clear separation between the two populations (see Supplementary Materials), demonstrating that ProClust can resolve fine-grained subpopulations when guided by lineage-informative features.

### 2.4 ProClust identifies biologically supported substructure across Tabula Muris tissues

To evaluate ProClust in biologically heterogeneous tissues, we applied it to five Tabula Muris FACS datasets comprising lung, pancreas, spleen, heart, and brain non-myeloid cells. These datasets differed in cellular composition, population abundance, and tissue complexity. A combined UMAP embedding showed the distribution of the reference cell types across the five datasets (Fig. 5a). We next focused on the brain non-myeloid and lung datasets to compare the clustering granularity produced by ProClust and existing methods and to determine whether the additional populations identified by ProClust represented transcriptionally distinct cellular subpopulations. Clustering results and performance comparisons for the pancreas, spleen, and heart datasets are provided in Supplementary Section S4.

**Figure 5:**
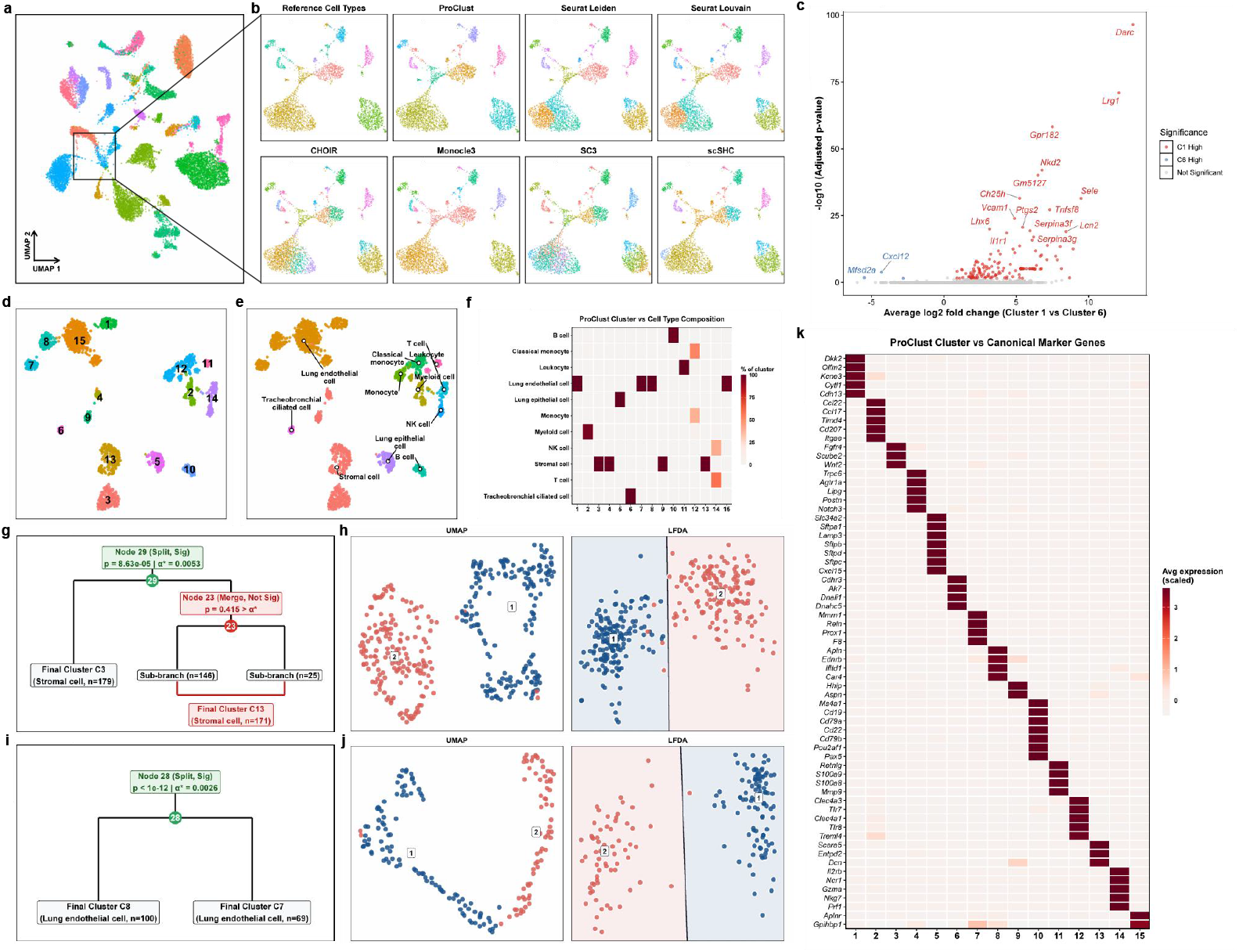
Comparison of clustering results and biological characterization of ProClust clusters in Tabula Muris datasets. **a**, UMAP embedding of cells from five Tabula Muris FACS datasets (lung, pancreas, spleen, heart, and brain non-myeloid), colored by reference cell-type annotation. The boxed region marks the brain non-myeloid cells examined in **b** and **c. b**, UMAP embeddings of brain non-myeloid cells colored by reference annotation or by clusters identified by ProClust, Seurat Leiden, Seurat Louvain, CHOIR, Monocle3, SC3, and scSHC. **c**, Volcano plot of differential expression between ProClust clusters 1 and 6 in the brain non-myeloid dataset. Both clusters correspond predominantly to cells annotated as endothelial in the reference. Negative average log_2_-fold changes indicate higher expression in cluster 6, whereas positive values indicate higher expression in cluster 1. Selected genes include *Mfsd2a* and *Cxcl12* in cluster 6 and *Darc, Sele, Lrg1*, and *Lcn2* in cluster 1. **d**, UMAP embedding of lung cells colored by the 15 ProClust clusters. **e**, The same embedding colored by reference cell-type annotation. **f**, Heatmap showing the reference cell-type composition of each ProClust lung cluster. Color indicates the percentage of cells in each cluster assigned to the indicated reference cell type. **g**, Portion of the ProClust hierarchical decision tree containing stromal nodes 29 and 23. The split at node 29 was retained (*p* = 8.63 × 10^*−*5^, *α*^*\**^ = 0.0053). The split at node 23 was not retained (*p* = 0.415), resulting in final clusters 3 and 13. **h**, UMAP and local Fisher discriminant analysis (LFDA) embeddings of the 350 cells at node 29, colored by the candidate groups. The line in the LFDA embedding indicates the classification boundary. **i**, Portion of the decision tree showing the retained split at lung endothelial node 28 (*p <* 10^*−*12^, *α*^*\**^ = 0.0026), resulting in final clusters 8 and 7. **j**, UMAP and LFDA embeddings of the 169 cells at node 28, colored by the candidate groups. The line in the LFDA embedding indicates the classification boundary. **k**, Heatmap showing scaled average expression of selected, literature-supported marker genes across the 15 ProClust lung clusters. The expression patterns support the biological interpretation of the clusters.

In the brain non-myeloid dataset, which contained seven reference-annotated cell types, the evaluated methods identified between 10 and 31 clusters (Fig. 5b). ProClust and Monocle3 each identified 10 clusters and achieved the two highest ARIs, at 0.8235 and 0.8199, respectively. The remaining methods identified between 12 and 31 clusters, with ARIs ranging from 0.28 to 0.73. In the lung dataset, which contained 11 reference-annotated cell types, ProClust identified 15 clusters (Fig. 5d,e) and Monocle3 identified 14. ProClust achieved an ARI of 0.53 and an NMI of 0.67, compared with 0.60 and 0.69 for Monocle3. The remaining methods identified between 16 and 28 clusters, with ARIs ranging from 0.29 to 0.49 and NMIs from 0.55 to 0.66. Although Monocle3 showed slightly higher agreement with the lung reference annotations, agreement alone does not establish whether distinct cellular populations have been appropriately retained. For example, Monocle3 assigned a subset of reference-annotated tracheobronchial ciliated columnar epithelial cells to a predominantly stromal cluster, whereas ProClust retained this small epithelial population as a separate cluster.

In the brain non-myeloid dataset, ProClust further separated the reference-annotated vascular endothelial cells into clusters 1 and 6. Differential expression analysis revealed distinct transcriptional profiles between the two clusters (Fig. 5c). Cluster 6 showed higher expression of *Mfsd2a, Cxcl12, Emcn, Nostrin*, and *Hmcn1*, consistent with a capillary-like endothelial phenotype. In contrast, cluster 1 was enriched for *Ackr1* (formerly *Darc*), *Sele, Lrg1, Lcn2*, and *Cd14*, consistent with a venous or inflammation-responsive endothelial phenotype. These expression differences support the interpretation of clusters 1 and 6 as transcriptionally distinct endothelial subpopulations.

The ProClust lung clusters largely corresponded to the reference cell types, while resolving additional populations within the endothelial and stromal compartments (Fig. 5f). Among the predominantly stromal cells, node 29 contained 350 cells and showed evidence against a single-population model (*p* = 8.633 × 10^*−*5^, *α*^*\**^ = 0.0053). Its split was retained, separating cluster 3 from the branch that ultimately formed cluster 13 (Fig. 5g,h). Within the latter branch, the split at node 23 was not significant (*p* = 0.415), and its two sub-branches were merged. A similar analysis of the predominantly endothelial cells retained the split at node 28 (*n* = 169; *p <* 10^*−*12^, *α*^*\**^ = 0.0026), yielding clusters 7 and 8 (Fig. 5i,j). The candidate groups at both nodes showed distinct distributions in the node-specific UMAP and LFDA representations. These node-level decisions provide statistical support for the retained subdivisions, which we further characterize using marker-gene expression below.

The 15 ProClust lung clusters showed distinct expression patterns of manually curated cell-type markers obtained from LungGENS and supported by the literature (Fig. 5k) (Du et al., 2021). The endothelial populations separated by ProClust showed different marker profiles: cluster 7 expressed *Prox1, Reln*, and *Mmrn1*, consistent with lymphatic endothelium (Du et al., 2021), whereas cluster 8 expressed *Car4, Ednrb*, and *Apln*, consistent with an aerocyte (aCap)-like phenotype. A third endothelial cluster, cluster 15, expressed *Aplnr* and *Gpihbp1*, consistent with a general capillary (gCap)-like phenotype (Gillich et al., 2020). The stromal clusters were also distinguishable by marker expression: cluster 3 expressed *Wnt2*, cluster 9 expressed the peribronchial fibroblast-associated genes *Hhip* and *Aspn* (Tsukui et al., 2020), and cluster 13 expressed *Scara5, Entpd2*, and *Dcn*. Other clusters likewise showed cell-type-associated expression patterns. Cluster 5 expressed the alveolar type II epithelial markers *Sftpa1, Sftpb, Sftpc*, and *Sftpd*, whereas cluster 6 expressed the cilia-associated genes *Ak7, Dnai1*, and *Dnahc5*, consistent with tracheobronchial ciliated columnar epithelial cells (Negretti et al., 2021). Among immune populations, cluster 10 expressed B-cell markers *Ms4a1, Cd19*, and *Cd79a*; cluster 11 expressed granulocyte-associated genes *Retnlg, S100a8, S100a9*, and *Mmp9* ; cluster 12 expressed dendritic-cell-associated genes *Clec4a3, Clec4a1*, and *Treml4* ; and cluster 14 expressed natural killer cell markers *Il2rb, Ncr1, Nkg7*, and *Prf1* (Negretti et al., 2021). These marker profiles provide biological context for the statistically retained endothelial and stromal subdivisions and support the interpretation of the other ProClust clusters.

### 2.5 proclust reveals transcriptional continuity across annotated cell-state boundaries in *Hydra*

In addition to assessing the statistical significance of clusters identified by a clustering algorithm, as demonstrated in the previous applications, ProClust can also be used to assess statistical support for distinctions among annotated cell types and states and to refine these annotations. To illustrate this capability, we apply ProClust to the reference annotations from the *Hydra* single-cell atlas (Siebert et al., 2019). The *Hydra* atlas identifies three major cell lineages: ectodermal epithelial (EC), endodermal epithelial (EN), and interstitial stem cell (IC) lineages (Fig. 6a). We applied ProClust and benchmark methods within these lineages, using the annotated cell types as the initial clustering structure (Fig. 6b). In particular, significance-based methods, including scSHC, CHOIR, and ProClust, were used to test whether distinctions between annotated states were statistically supported under their respective testing frameworks. By contrast, Monocle3, Louvain, and Leiden do not provide statistical significance measures for annotated cell states; for these methods, the reference annotations were used only to initialize the clustering procedure.

**Figure 6:**
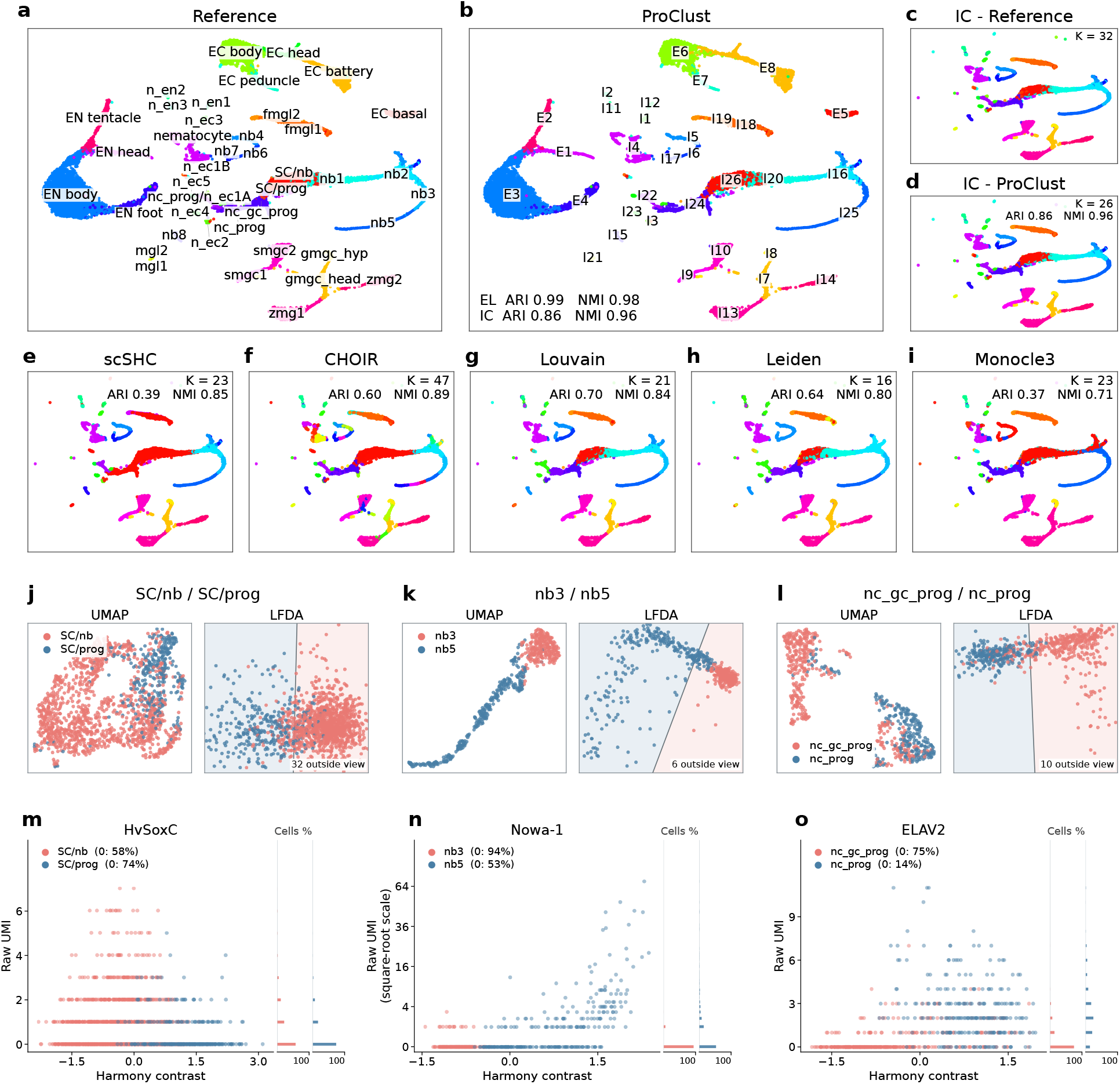
Comparison of clustering results and biological validation in the *Hydra* single-cell atlas. **a–b**, UMAP embeddings of *Hydra* cells colored by the reference cell-type annotations and ProClust clusters, respectively. **c–i**, UMAP embeddings of interstitial stem-cell populations in *Hydra*, colored by the reference annotations and clustering results from the benchmark methods. ProClust and all benchmark methods were initialized using the reference annotations. **j–l**, UMAP and LFDA embeddings of cells belonging to ProClust clusters with *p*-values above the adjusted significance threshold of 0.05. For each cell-type pair – SC/nb versus SC/prog, nb3 versus nb5, and nc_gc_prog versus nc_prog – normalization and Harmony-PCA dimensionality reduction were performed separately within the corresponding pair, and both UMAP and LFDA embeddings were computed from the resulting Harmony PCs. **m–o**, Distribution of marker-gene expression for the cells shown in **j–l** as a function of the Harmony contrast between the corresponding annotated cell types. A distinct ProClust cluster is expected to exhibit a separate expression mode together with separation along the Harmony contrast, rather than being concentrated within the overlapping region between the annotated cell types.

Compared with existing methods, ProClust achieves substantially better agreement with the reference annotations in the IC lineage (Fig. 6c-d). It attains an ARI of 0.86 while merging putative clusters and reducing the number of clusters from 32 to 26. In contrast, scSHC and CHOIR tend to either over-merge or over-split the reference annotations (Fig. 6e-f), resulting in lower ARIs of 0.39 and 0.60, respectively. Furthermore, conventional clustering algorithms such as Louvain, Leiden, and Monocle3 do not provide statistical significance measures for determining whether existing annotations should be merged or split. Consequently, their resulting clusters do not necessarily preserve the boundaries of the reference annotations and may introduce broader re-partitioning beyond simple merging or splitting. Therefore, these methods can further yield new putative clusters that require additional biological re-annotation (Fig. 6g-i). They also exhibit lower ARI and NMI than ProClust. For the EC and EN lineages, ProClust achieves performance comparable to that of the other methods, with detailed results provided in the Supplementary Materials.

ProClust suggests merging SC/nb with SC/prog (stem/progenitor, *n* = 1,546), nb3 with nb5 (nematoblast, *n* = 541), and nc_gc_prog with nc_prog (neuronal progenitor, *n* = 648). All three pairs have unadjusted *p*-values exceeding 0.5, providing little statistical evidence for separated cluster structure (Fig. 6j-l). For SC/nb and SC/prog in particular, the LFDA embedding does not reveal two separable clusters, consistent with the corresponding UMAP visualization. Moreover, the boundaries of the annotated cell states do not align with visually distinct regions in the UMAP embedding. For the nb and nc pairs, LFDA reveals elongated or heavy-tailed structures rather than clearly separated clusters, whereas the existing annotations divide cells within relatively concentrated regions of the embedding. Thus, the ambiguity of these annotations extends beyond whether a statistically detectable mixture structure exists – the current annotation boundaries may fail to correspond to the dominant structure in the data. The UMAP embedding of the nb cells is consistent with this interpretation, showing only weak separation between the annotated states. For the nc cells, UMAP appears to produce two visually distinct groups; however, this separation is not evident in the corresponding LFDA, PCA, or t-SNE (Van der Maaten and Hinton, 2008) representations and may therefore reflect a visualization artifact rather than robust cluster structure (see Supplementary Materials). Because nonlinear embeddings such as UMAP can introduce or exaggerate apparent separation, we use them only as qualitative visual references.

Biologically, the annotations of SC, nb, and nc states are associated with the expression of *HvSoxC, Nowa-1*, and *ELAV2*, respectively (Siebert et al., 2019; see Supplementary Materials). These annotations arise within the highly dynamic interstitial lineage of *Hydra*, in which multipotent interstitial stem cells continuously generate neuronal, nematocyte, and gland-cell progeny through intermediate differentiation states (David, 2012; Wenger et al., 2016; Holstein, 2023). Expression of the three marker genes is strongly associated with the corresponding annotation contrasts (Fig. 6m-o). However, their known biological roles suggest that they primarily reflect differentiation programs or maturation stages rather than defining individual states. For example, *HvSoxC* is expressed in transitional interstitial cells between candidate stem cells and differentiating neuronal and nematoblast lineages, and has been interpreted as a marker of cells undergoing differentiation (Siebert et al., 2019; Sprecher, 2022). Consistent with this broader role, we observe *HvSoxC* expression in both SC/nb and SC/prog, with a higher detection frequency in annotated SC/nb cells. For nb states, more recent analyses place nb3 and nb4 among the earliest post-mitotic nematoblast states, whereas nb5–nb8 correspond to capsule-forming stages in which *Nowa-1* and other nematocyst-associated genes become expressed (Moneer et al., 2021). Finally, *ELAV2* is also broadly expressed across the neuronal lineage, including neuronal progenitors and differentiated neurons (Little et al., 2025).

Importantly, within each of these candidate cell-state pairs, the corresponding marker expression distributions remain dominated by a mode at zero and do not exhibit a clear secondary expression mode (Fig. 6m-o). In particular, *Nowa-1* expression shows no significant deviation from a negative binomial distribution (Pearson chi-square test *p* = 0.395). Thus, although marker expression differs across the annotated groups, differential expression alone does not establish that the annotation boundary corresponds to two biological populations. Such differences may instead reflect changes along a continuous differentiation trajectory. Together with the lack of significant clustering structure identified by ProClust, these results suggest that the current annotations capture biologically meaningful differentiation signals, but their precise boundaries may not correspond to separable cell states for all cells.

## 3 Discussion

An important challenge in single-cell transcriptomic analysis is determining whether a clustering result contains too few, too many, or an appropriate number of clusters. Commonly used clustering methods can produce different results depending on analytical choices, but generally provide limited statistical evidence for determining whether candidate clusters should remain separate or be merged. As a result, both overclustering and underclustering may occur without a clear statistical basis for selecting the final clustering result. ProClust addresses this problem by testing statistical significance of derived clusters. The resulting *p*-values can inform researchers regarding which candidate divisions should be retained and which should be merged. Across the settings examined in this study, ProClust reduced unsupported splitting while preserving biologically meaningful subpopulations.

The simulation results highlight the value of using statistical evidence as a stopping rule for clustering refinement. Conventional clustering algorithms will return a partition even when the data arise from a single population, making it difficult to distinguish meaningful structure from divisions created by the clustering procedure itself. Under the null hypothesis, ProClust avoided retaining such unsupported divisions in both simulation frameworks. Its lower-tail behavior is useful in this setting because false subdivisions may subsequently be interpreted as distinct cell types or states. Importantly, this protection against overclustering did not result in systematic merging of genuine populations. Under the alternative hypothesis, ProClust generally recovered both the correct number of populations and assignments closely aligned with the true labels, including in the PBMC-based simulations that preserved features of real single-cell expression data. Taken together, these findings suggest that ProClust provides a practical balance between avoiding unsupported divisions and retaining population structure supported by the data.

The real-data analyses illustrate the complementary roles of statistical and biological evidence in defining cellular populations. Reference annotations provide biological context through established markers and functional knowledge, whereas statistical testing assesses whether a candidate division is supported by the observed expression structure. ProClust contributes to this process by retaining subdivisions supported by the data rather than accepting every division generated by the initial clustering. A subdivision is particularly compelling when statistical support is accompanied by coherent marker expression or known functional differences. Differences between the inferred structure and existing annotations can also be informative, as they may point to cell states or subpopulations that were not distinguished in the original annotation. In the HLCA, Tabula Muris, and *Hydra* analyses, ProClust showed strong agreement with established annotations while preserving additional subdivisions with interpretable marker-expression patterns. ProClust can therefore complement biological annotation by providing statistical evidence for selecting clustering granularity and identifying populations that merit further investigation.

ProClust also integrates statistical testing with useful visualization of candidate clusters. Non-linear embeddings such as UMAP are useful for summarizing the overall structure of a dataset, but they are not designed to provide statistical evidence for individual clustering decisions. Differences between closely related populations may also be obscured when the embedding is dominated by variation among major cell classes. ProClust addresses this issue by constructing a two-dimensional LFDA representation separately for each tested node. This linear representation focuses on variation relevant to the candidate partition. Consequently, the projected observed data and synthetic samples from the fitted single-population null model can be evaluated within the same coordinate system and using the same fixed decision regions. Learning a projection that emphasizes candidate cluster differences can, however, make separation appear stronger than it would in independent data. Cross-fitting is therefore essential because it separates the observations used to learn the LFDA projection from those used to evaluate statistical significance. The combination of a locally informative representation and held-out evaluation allows ProClust to improve the interpretability of clustering decisions without treating visual separation alone as evidence of distinct populations.

Like any clustering refinement and testing procedure, the conclusions produced by ProClust depend on the input clusters. The initial clustering must therefore be sufficiently informative, because ProClust can merge existing base clusters but cannot recover distinct populations that have already been mis-classified by a clustering algorithm. The hierarchy constructed from these base clusters imposes an additional constraint by determining which groups can be considered for merging. If related base clusters are placed in different upper-level branches and the intervening split is retained, they cannot be reconsidered together later in the procedure. ProClust further relies on appropriate quality control, normalization, and feature-screening procedures for scRNA-seq data. Biologically meaningful differences may be missed when they are not captured by the selected features. Conversely, outlying cells may distort the locally fitted LFDA representation, while technical variation may create separation that does not correspond to the biological structure of interest. Appropriate quality control, feature selection, batch correction, and robust handling of outliers therefore remain important.

These limitations suggest several directions for further development of ProClust. Future work could relax the fixed-tree constraint by developing procedures that allow related base clusters assigned to different branches of the initial hierarchy to be reconsidered for merging. Such an extension would require corresponding adjustments to the testing framework to account for the additional data-driven selection. Future work could also explore more robust approaches to handling outlying cells during node-specific projection and systematically evaluate how feature selection affects ProClust’s ability to detect subtle population differences. Despite these limitations, our results show that ProClust provides a useful statistical framework for refining an initially overclustered solution. By providing statistical evidence and node-specific LFDA visualizations for decisions about whether candidate divisions should be retained or merged, ProClust offers a more transparent basis for selecting the final clustering granularity and identifying candidate subpopulations for further biological investigation.

## 4 Methods

### 4.1 ProClust Algorithm

ProClust takes as input an original count matrix **X**, an initial clustering partition into *K* base clusters, and a low-dimensional representation **Y** of the cells (typically PCA or a batch-corrected embedding computed upstream). It builds a hierarchical tree over the *K* base clusters and sequentially tests every internal node of that tree for genuine subpopulation structure, using a cross-validated, projection-based statistic at each node (Liu et al., 2008; Kimes et al., 2017).

#### Cluster tree construction

The base clusters are organized into a binary tree that determines the order in which internal nodes are tested and which clusters are merged at each node. By default, ProClust builds this tree from a graph-based connectivity measure between clusters while allowing for user-specific hierarchical clusters. **Y** is truncated to its first *d* columns (default *d* = 50, or fewer if **Y** has fewer columns) and column-standardized; a *k*-nearest-neighbor graph is built over cells in this space, with neighborhood size 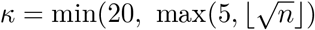, *n* − 1), where *n* is the total number of cells, and converted to a Jaccard-weighted shared-nearest-neighbor graph following standard Seurat preprocessing (Stuart et al., 2019). For every pair of base clusters *i, j* we compute the mean graph edge weight between all cells in cluster *i* and all cells in cluster *j*, giving a *K* × *K* inter-cluster connectivity matrix **C** with self-connectivity fixed at 1 on the diagonal. Each cluster’s row of **C** is taken as its connectivity profile to every other cluster, and *D*_*ij*_ = ∥*C*_*i,·*_ − *C*_*j,·*_∥_2_ standard agglomerative clustering (complete linkage) is applied to **D**. This tree-construction step is modular: the sequential testing procedure below depends only on the resulting merge order, heights, and leaf structure, so the tree can be built by other methods without changing any other part of ProClust.

#### Sequential testing with node-specific error control

Given the tree, each internal node *v* (there are *K* − 1 of them) is associated with a leaf set, the base clusters merged at or below *v*, and with the cells whose base label falls in that leaf set. Internal nodes are tested top-down, in decreasing order of merge height, so the root, spanning all *K* base clusters, is tested first. Testing is sequential and closed: a node is tested only if all of its ancestors were classified significant, and testing along a branch halts as soon as a node is classified not significant.

To adjust the *p*-values for multiple testing, each tested node *v* is assigned its own significance threshold 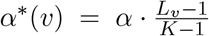, where *L*_*v*_ is the number of base clusters merged at or below *v* (Kimes et al., 2017). The root (*L*_*v*_ = *K*) is tested at level *α*; a node merging two base clusters is tested at 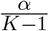. Nodes with fewer than *n*_min_ cells are not tested. A node whose test returns an error is, by default, classified significant; the alternative behavior classifies it not significant.

At each tested node, the count matrix restricted to that node’s cells is renormalized (log-normalization, scaling across all genes, and PCA recomputed locally rather than reused from *Y*). The number of PCs computed at a node is 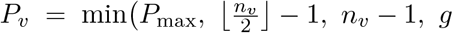, [2*L*_*v*_ if *n*_*v*_ *<* 500], where *n*_*v*_ is the node’s sample size, *g* is the number of genes, and *P*_max_ is a fixed hyperparameter (default 50) capping the number of PCs computed at any node. If a node contains more than one batch, batch integration (Korsunsky et al., 2019) is applied to this local PCA embedding. The node’s local sub-tree (the master tree restricted to the base clusters present at that node) is carried forward for use in the training-label initialization step below.

#### Node-level test

At each node, ProClust tests whether the observed cluster separation is stronger than would be expected under a single log-concave population. The procedure consists of the following steps:

1. Randomly split the cells’ node-specific PC embedding into a training set, **Y**_tr_, and a testing set, **Y**_te_.
2. Apply a clustering algorithm to the training data to obtain cluster labels 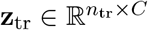, *C* ∈ {2, 3}.
3. Using **Y**_tr_ and **z**_tr_, fit a LFDA projection 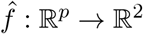.
4. Apply the learned projection to the testing data and compute the observed cluster index, CI_**Y**_, from the two-dimensional representation 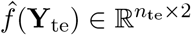 and 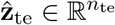.
5. Generate *B* null datasets from a fitted log-concave distribution based on 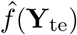, and compute the corresponding null cluster indices CI_1_, …, CI_*B*_.
6. The node-level *p*-value is obtained by approximating the null distribution of the cluster index with a normal distribution.

To improve data efficiency and reduce variability due to sample splitting, the procedure is repeated over *R* independent outer restarts, with *R* = 20 by default. Details of the construction of the training labels, the LFDA projection, the cluster index, and the generation of log-concave null datasets are provided in the following sections.

#### Construction of training labels

If the node’s base labels resolve to more classes than a target ceiling *C* (default *C* = 3), they are collapsed by cutting the node’s local sub-tree at *C* groups. A *d*-dimensional, column-standardized representation of the training cells is built from their top local PCs, and *k*-means with *C* centers is run on this space. When the collapsed labels give usable (finite, distinct) group centroids, *k*-means is warm-started from them with a single run; when no usable guide labels are available at all, *k*-means is instead run unguided with ten random restarts.

#### LFDA embedding

LFDA (target dimensionality 2) is fit on the training cells using the training labels as classes, attempted on a descending ladder of input dimensionalities, { *p*_avail_, 50, 30, 20, 10, 5, 2 }∩ {2, …, *p*_avail_}, where *p*_avail_ is the number of available local PCs; the first dimensionality at which the fit succeeds is used, with the neighborhood size in its affinity graph set to *κ*_LFDA_ = max(1, min(7, *n*_class,min_ − 1)), where *n*_class,min_ is the size of the smallest class among the training cells in this fold.

#### Cluster index

The cluster index (Liu et al., 2008; Keefe and Marron, 2025) is computed on the LFDA embeddings 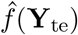 and their corresponding predicted labels 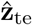, defined as

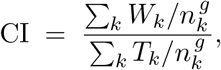

where, for cluster *k* of size *n*_*k*_, *W*_*k*_ is the sum, over cells in *k* and over embedding axes, of the absolute deviation of each cell from its own cluster’s mean raised to power *l*, and *T*_*k*_ is the analogous sum of absolute deviations from the global mean computed over the same cells. The default is *l* = 1 and *g* = 0.5. A smaller CI indicates a higher level of separation among clusters, suggesting genuine clustering structure.

#### Null log-concave model

The null distribution can be estimated and sampled directly using the R package LogConDEAD (Cule et al., 2010). However, this approach can be computationally intensive for large scRNA-seq datasets. As a more efficient alternative, ProClust fits a bivariate null distribution directly to 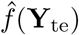. It considers three candidate families of log-concave distributions – the skewed generalized error distribution (SGED) (Theodossiou, 2015), Beta distributions, and uniform distributions. The candidate providing the best fit to the projected data is selected to generate null samples. These distributions span the full range of cluster index values attainable under log-concave distributions. For the SGED candidate, each axis is fit marginally with

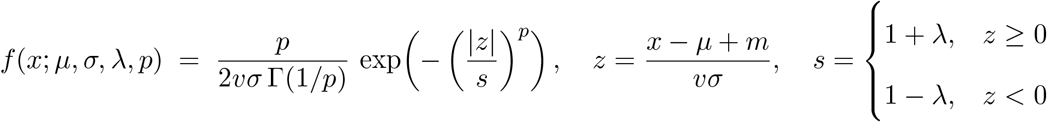

with position and scale terms

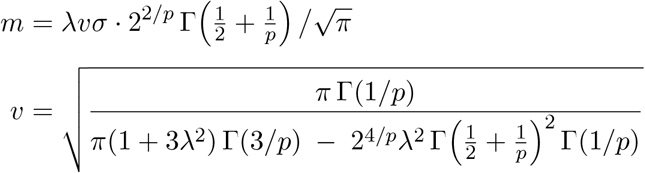

reducing to a symmetric generalized error distribution at *λ* = 0 and to a Gaussian at *λ* = 0, *p* = 2. Parameters are estimated by a moment-based rule, solving 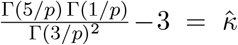 for *p* via root-finding, where 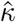 is the sample excess kurtosis, and 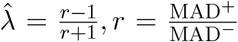 . The other two candidates are a four-parameter rescaled Beta,

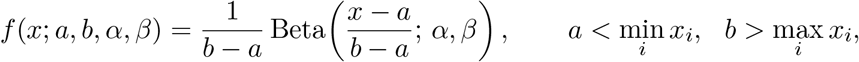

with support padded beyond the observed range and (*a, b, α, β*) fit by maximum likelihood, and a two-parameter uniform Unif(min_*i*_ *x*_*i*_, max_*i*_ *x*_*i*_). For all candidates, the two fitted marginal CDFs 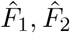 are coupled with a bivariate Gaussian copula, with correlation 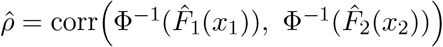, shrunk to [−0.98, 0.98]. If the selected fit yields a non-finite copula transform 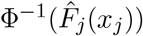 on either axis, the selection is discarded and both axes are refit as SGED.

#### Monte Carlo evaluation of the null

ProClust generates *B* synthetic two-dimensional datasets from the fitted null distribution. Each synthetic dataset is passed through the same LFDA classifier to obtain predicted cluster labels 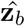, *b* = 1, …, *B*. The cluster index is then computed for each synthetic dataset, producing null statistics CI_1_, …, CI_*B*_. For each Monte Carlo replicate *b*, the signed difference Δ_*b*_ = CI_**Y**_ − CI_*b*_, is computed. The normalized *p*-value is computed as

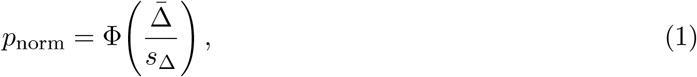

where 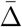 and *s*_Δ_ denote the mean and standard deviation of Δ_1_, …, Δ_*B*_, respectively. Within one outer restart, the signed differences from all two folds are pooled position-by-position, 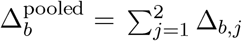, where only replicate indices for which both folds yield finite values are retained. Eq. (1) is then applied to 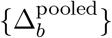 to give one *p*-value per restart. Finally, at the tree level, a node’s decision compares the unadjusted median of its *R* restart-specific *p*-values against *α*^*\**^(*v*).

#### Initial over-clustered partition

To provide a common input for the post-clustering merging and significance-testing methods evaluated below, we generated an over-clustered partition using either Leiden (Traag et al., 2019) or Louvain (Blondel et al., 2008), depending on the dataset. Monocle3 (Cao et al., 2019) was also included as a widely used graph-based clustering approach.

#### Competing post-clustering methods

We compared ProClust with two existing hierarchical frameworks that identify cluster structure through statistical testing and pruning: CHOIR (Sant et al., 2025) and scSHC (Grabski et al., 2023). scSHC was applied directly to the Leiden or Louvain partition described above. CHOIR constructs its own initial over-clustering internally and was therefore run independently. Detailed implementation procedures, preprocessing choices, and method-specific hyperparameter settings are provided in the Supplementary Methods.

#### Performance measures

Under *H*_0_, statistical calibration was assessed using the empirical distribution of the resulting *p*-values. A well-calibrated procedure is expected to produce sub-uniform *p*-values under the null hypothesis. Under *H*_1_, agreement between the estimated clustering and the true population labels was evaluated using the adjusted Rand index (ARI) and normalized mutual information (NMI). We additionally recorded the estimated number of clusters, 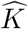, and compared it with the true number of populations, *K*_true_, in simulations or the number of reference populations, *K*_ref_, in real datasets. Computational runtime was recorded in seconds to assess the relative computational cost of each method. For the parametric simulations, the purity of the initial overclustered partition was also calculated as

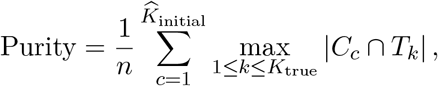

where *C*_*c*_ denotes an initial cluster and *T*_*k*_ denotes a true population.

### 4.2 Simulation study design

We used two distinct simulation frameworks to assess type I error control and clustering performance. In the parametric framework, each dataset contained 5,000 cells and 2,000 genes, with counts generated from negative-binomial marginal distributions incorporating gene–gene dependence and continuous latent-factor variation within each population. Under *H*_0_, all cells were generated from a single population with no population-specific expression shifts. Under *H*_1_, we considered *K*_true_ ∈ 3, 5, 7, 9 approximately equally sized populations, each defined by expression shifts in a nonoverlapping set of marker genes. In the PBMC-based semi-synthetic framework, scDesign2 was fitted separately to nine immune cell populations identified in the PBMC 3k reference dataset, thereby preserving population-specific expression distributions, overdispersion, and gene–gene dependence. Under *H*_0_, cells were generated exclusively from the largest reference population, which consisted predominantly of CD4^+^ T cells. Under *H*_1_, cells were generated from the *K*_true_ ∈ 3, 5, 7, 9 most abundant reference populations in proportion to their relative frequencies in the reference dataset. Each simulated dataset contained 5,000 cells, and 100 independent datasets were generated under each setting. Complete data-generating mechanisms and parameter specifications are provided in the Supplementary Methods.

### 4.3 Transcriptomic Datasets

#### Human Lung Cell Atlas

We benchmarked on the Human Lung Cell Atlas (HLCA) core dataset, comprising 584,944 cells and 27,402 genes from healthy lung tissue across 107 donors, with a five-level hierarchical cell type annotation (Sikkema et al., 2023). At the coarsest level (`ann_level_1`), cells fall into four compartments: Epithelial (282,065), Immune (229,496), Endothelial (48,166), and Stroma (25,217); annotation resolution increases through `ann_level_2–4` (11, 25, and 39 categories, respectively). We used `ann_level_4` as the reference annotation, since `ann_level_5` is defined for only a subset of cells (418,277 unassigned). Immune-compartment cells lacking an `ann_level_4` label were removed, and 10% of cells from each remaining level-4 immune subtype were randomly sampled (stratified, fixed seed), yielding a final benchmark set of 22,280 cells.

#### Tabula Muris

We analyzed five tissue-specific scRNA-seq datasets from the fluorescence-activated cell sorting (FACS) Smart-seq2 arm of the Tabula Muris atlas (Tabula Muris Consortium, 2018b): brain non-myeloid, heart, lung, pancreas, and spleen. Raw count matrices and cell-level metadata were obtained from the corresponding tissue-specific Seurat objects available through the Figshare repository (Tabula Muris Consortium, 2018a). Published cell-type annotations in the cell_ontology_class metadata field were used as reference labels, and cells without an annotation were removed. After filtering, the five datasets contained 12,675 cells in total, with individual dataset sizes ranging from 1,564 to 4,337 cells and the number of annotated cell types ranging from 3 to 11. Each dataset contained expression measurements for 23,341 genes.

#### *Hydra* stem cell transcriptomes

We analyzed three annotated subsets of the adult *Hydra vulgaris* Drop-seq atlas (Siebert et al., 2019): epithelial cells (EL; 6,742 cells and 9 annotated states), interstitial cells (IC; 7,797 cells and 32 annotated states), and neurons and neuronal progenitors (NN; 1,136 cells and 15 annotated states). EL combined 2,520 ectodermal and 4,222 endodermal epithelial cells. NN was entirely contained within IC, yielding 14,539 unique cells across the three analysis subsets. The transcriptome-based UMI count matrix contained 37,114 transcript features; excluding features with zero counts within each subset retained 31,564, 30,121, and 20,025 features for EL, IC, and NN, respectively. All three subsets contained cells from the same 10 whole-animal Drop-seq libraries. Library identity was used for Harmony batch adjustment.

### 4.4 Analysis configuration

#### Preprocessing and initial clustering

In the parametric and PBMC-based simulations and in the HLCA and Tabula Muris analyses, counts were normalized to a total expression of 10,000 per cell and log-transformed by using Seurat. All 2,000 genes were retained in the parametric simulations. For the PBMC-based simulations and each Tabula Muris dataset, the 2,000 most highly variable genes were selected using the Seurat variance-stabilizing transformation method. For HLCA, 2,000 highly variable genes were selected while accounting for dataset-specific effects. The resulting expression matrices were scaled and subjected to PCA, retaining 50 principal components for the simulated and HLCA datasets and 30 principal components for the Tabula Muris datasets. The five Tabula Muris tissues were processed independently, without explicit correction for animal or sequencing-plate effects or additional outlier filtering.

The parametric simulations, PBMC-based simulations, and Tabula Muris datasets used the same initial clustering procedure. A shared nearest-neighbor graph was constructed from the first 30 principal components, followed by Louvain clustering at a resolution of 1.2. The resulting high-resolution partition was used to initialize ProClust and scSHC.

For HLCA, Harmony was applied to the PCA representation using dataset identity as the batch variable, after which cells identified as potential outliers by the Local Outlier Factor algorithm were removed. A neighborhood graph was constructed from the Harmony-corrected representation using 30 nearest neighbors and cosine distance. Leiden clustering at a resolution of 1.0 was then used to obtain the initial partition for ProClust and scSHC.

#### Hyper-parameter configuration

ProClust used a consistent hyper-parameter specification throughout this paper. Node-level tests compared at most three candidate subclusters and used up to 30 or 50 principal components, 20 random initializations, two-fold cross-fitting, and 100 or 200 null replicates.

For the simulation analyses, batch correction was not applied because batch effects were not included in the data-generating mechanisms. For the Tabula Muris analyses, each tissue was analyzed separately without batch correction because preliminary examination indicated that batch-associated structure was limited. For the initial overclustered partitions supplied to ProClust and scSHC, Louvain clustering was performed at a resolution of 1.2, rather than the default value of 0.8, to generate sufficiently fine-grained starting partitions for subsequent significance-based merging. The corresponding shared nearest-neighbor (SNN) graphs were constructed using *k* = 10 nearest neighbors in the simulations to retain finer local structure and promote initial overclustering, whereas the default value of *k* = 20 was used for the Tabula Muris datasets. As independent benchmark methods, Seurat Louvain and Leiden were run at the default clustering resolution of 0.8. They used the same graph-construction settings as the corresponding initial partitions, with *k* = 10 in the simulations and *k* = 20 in the Tabula Muris analyses, to avoid differences attributable solely to the underlying SNN graph. For CHOIR, the pruning significance threshold was set to its default value of *α* = 0.05. scSHC used 2,000 features, rather than its default maximum of 2,500, to match the feature-selection scale of the common preprocessing workflow, while retaining its default setting of 30 principal components. Monocle3 retained 30 principal components during preprocessing to match the dimensionality used in the common preprocessing workflow. UMAP dimensionality reduction and the default Leiden clustering algorithm were then applied with an explicitly specified resolution of 10^*−*5^, following a documented Monocle3 analysis configuration rather than the automatic default setting of NULL. For SC3, the full raw-count matrix was supplied as input, and the number of clusters was estimated using `sc3_estimate_k` to avoid specifying it using the ground-truth labels. Clustering was then performed using SC3’s default gene-filtering procedure.

For HLCA, the initial Leiden partition used by ProClust was obtained at a resolution of 1.0. ProClust’s hierarchical tree over the initial Leiden clusters was constructed by computing the centroid of each cluster in the Harmony-corrected PCA embedding and applying Ward’s hierarchical clustering (ward.D2 linkage) to the Euclidean distances between cluster centroids. CHOIR used Harmony batch correction with dataset identity as the batch variable and a default pruning significance threshold of *α* = 0.05. To ensure a fair comparison with ProClust, scSHC was aligned to use the same input dimensions of 2000 features and 50 principle components, with dataset identity specified as the batch variable.

