## Supplementary Information for "ProClust provides reliable statistical significance of clusters for single-cell transcriptomic data"

### S1 Supplementary Methods

We considered six competing clustering methods: Leiden, Louvain, CHOIR, scSHC, Monocle3 and SC3. The implementation details and method-specific parameters used in our analyses are described below.

**Leiden** Leiden clustering (Traag et al., 2019) was performed on a $k$-nearest-neighbor graph constructed from the batch-corrected embedding. The clustering resolution controlled the granularity of the resulting partition, while the neighborhood size and distance metric were specified during construction of the neighborhood graph.

**Louvain** Louvain clustering (Blondel et al., 2008) was performed on a shared nearest-neighbor graph constructed from the same batch-corrected embedding. The resolution parameter controlled the granularity of the resulting clustering, with the neighborhood graph construction held consistent with the other graph-based methods where applicable.

**CHOIR** CHOIR (Sant et al., 2025) was applied using its two-step procedure consisting of hierarchical tree construction with `buildTree` followed by statistical pruning with `pruneTree`. The significance threshold $\alpha$ used during pruning determines whether candidate splits in the hierarchical tree are retained as distinct clusters.

**scSHC** scSHC (Grabski et al., 2023) was applied using its hierarchical significance-testing procedure implemented in `testClusters`. The number of features (`num_features`), the number of principal components (`num_PCs`) used for dimensionality reduction, and the significance threshold $\alpha$ were specified as the primary method-specific parameters governing the hierarchical testing procedure.

**Monocle3** Monocle3 clustering (Cao et al., 2019) was performed using `cluster_cells`. The neighborhood size $k$ used to construct the nearest-neighbor graph and the underlying clustering algorithm (by default, the Leiden algorithm) serve as the primary drivers of cell grouping, with larger neighborhood sizes generally producing coarser graph representations.

**SC3** SC3 (Kiselev et al., 2017) was applied using its consensus clustering framework, which combines the results of multiple $k$-means runs. The number of clusters $k$, estimated using `sc3_estimate_k`, and the gene-filtering procedure were the primary method-specific parameters governing the resulting partition.

### S2 Simulation Data-Generating Mechanisms

#### S2.1 Parametric simulation

Each simulated dataset contained 5,000 cells and 2,000 genes. Under $H_0$, all cells were generated from a single population without population-specific expression differences. Under $H_1$, we considered $K_{\text{true}} \in \{3, 5, 7, 9\}$ populations, with cells allocated as evenly as possible across populations. One hundred independent datasets were generated for each setting.

For each dataset, the gene-specific baseline log-expression means were independently sampled from a normal distribution with mean zero and standard deviation 1. Under $H_1$, each population was assigned a distinct set of 20 nonoverlapping marker genes. The log-expression mean of each marker gene was increased by 0.5 in its corresponding population. No population-specific expression shifts were introduced under $H_0$.

To introduce continuous within-population heterogeneity, three latent factors were generated for each population. Each latent factor affected a randomly selected 10% of the genes through standardized normal loadings. Cell-specific latent scores were sampled independently from a three-dimensional standard normal distribution. The resulting latent-factor effects were scaled by 0.4 and added to the log-expression means.

Cell-specific library sizes were sampled from a log-normal distribution with a median of 20,000 and a log-scale standard deviation of 0.2. The gene-specific means for each cell were subsequently rescaled so that their sum matched the sampled library size.

Gene–gene dependence was introduced using a Gaussian copula with an autoregressive correlation structure,

$$\Sigma_{gg'} = 0.2^{|g-g'|}, \tag{S1}$$

where $\Sigma_{gg'}$ denotes the correlation between genes $g$ and $g'$. Counts were then generated from negative-binomial marginal distributions with dispersion parameter $\theta = 10$. No additional zero inflation or batch effects were included.

#### S2.2 PBMC-based semi-synthetic simulation

To complement the controlled parametric simulations, we constructed a reference-based semi-synthetic simulation using the 10x Genomics peripheral blood mononuclear cell dataset (PBMC 3k). The reference dataset contained approximately 2,700 cells. Raw counts were processed using Seurat, and the 2,000 most highly variable genes were selected using the variance-stabilizing transformation method. Graph-based clustering with a resolution of 0.8 identified nine reference populations representing major immune cell types. These populations were ranked according to their observed abundances in the reference dataset for subsequent data generation.

The scDesign2 framework was fitted separately to each of the nine reference populations using the 2,000 selected genes. For each population, scDesign2 estimated gene-specific negative-binomial marginal distributions and a Gaussian copula dependence structure. This procedure preserved the population-specific expression levels, overdispersion, and gene–gene dependence observed in the reference data.

Under $H_0$, 5,000 cells were generated exclusively from the largest reference population, which consisted predominantly of $CD4^+$ T cells. This produced a single-population dataset with realistic within-population variability but no true discrete population structure.

Under $H_1$, we considered $K_{\text{true}} \in \{3, 5, 7, 9\}$. For each value of $K_{\text{true}}$, the $K_{\text{true}}$ most abundant reference populations were selected. Their observed relative frequencies were renormalized across the selected populations and used to allocate a total of 5,000 simulated cells. One hundred independent datasets were generated for each setting.

### S3 Additional Simulation Results

#### S3.1 Complete benchmark results

Complete benchmark results are reported in Supplementary Tables S1 and S2. Values are mean ± standard deviation across 100 replicates per setting.

SC3 was run with automatic cluster-number estimation enabled (`k_estimator = TRUE`). In the parametric simulation with $K_{\text{true}} = 5, 7, 9$, it returned a single cluster in all replicates, yielding zero ARI and NMI. These single-cluster estimates bypassed subsequent consensus clustering, resulting in substantially shorter runtimes.

Table S1: **Clustering performance in the parametric simulation.** The ground-truth number of clusters was $K_{\text{true}} = 3, 5, 7$, or 9. Values are mean ± standard deviation across 100 replicates per setting. Runtime is reported in seconds.

| Setting | Method | ARI | NMI | $\widehat{K}$ | Runtime (s) |
|---|---|---|---|---|---|
| $K_{\text{true}} = 3$ | ProClust | $0.998 \pm 0.001$ | $0.996 \pm 0.003$ | $3.00 \pm 0.00$ | $285.7 \pm 12.6$ |
| | CHOIR | $0.430 \pm 0.118$ | $0.485 \pm 0.057$ | $11.51 \pm 2.12$ | $4560.2 \pm 1105.5$ |
| | scSHC | $0.533 \pm 0.303$ | $0.550 \pm 0.285$ | $4.20 \pm 2.32$ | $725.2 \pm 337.4$ |
| | Monocle3 | $0.999 \pm 0.001$ | $0.997 \pm 0.002$ | $3.00 \pm 0.00$ | $26.7 \pm 2.9$ |
| | SC3 | $0.443 \pm 0.062$ | $0.520 \pm 0.036$ | $8.58 \pm 1.06$ | $6464.9 \pm 690.4$ |
| $K_{\text{true}} = 5$ | ProClust | $0.996 \pm 0.002$ | $0.993 \pm 0.003$ | $5.00 \pm 0.00$ | $408.5 \pm 28.3$ |
| | CHOIR | $0.631 \pm 0.113$ | $0.666 \pm 0.064$ | $13.34 \pm 2.70$ | $3786.5 \pm 857.9$ |
| | scSHC | $0.548 \pm 0.264$ | $0.603 \pm 0.249$ | $7.88 \pm 3.67$ | $1045.7 \pm 359.4$ |
| | Monocle3 | $0.997 \pm 0.001$ | $0.994 \pm 0.003$ | $5.00 \pm 0.00$ | $23.2 \pm 2.6$ |
| | SC3 | $0.000 \pm 0.000$ | $0.000 \pm 0.000$ | $1.00 \pm 0.00$ | $77.7 \pm 11.1$ |
| $K_{\text{true}} = 7$ | ProClust | $0.995 \pm 0.002$ | $0.992 \pm 0.003$ | $7.00 \pm 0.00$ | $491.8 \pm 47.6$ |
| | CHOIR | $0.716 \pm 0.140$ | $0.766 \pm 0.090$ | $12.35 \pm 3.52$ | $4256.4 \pm 1350.3$ |
| | scSHC | $0.603 \pm 0.314$ | $0.640 \pm 0.311$ | $9.51 \pm 4.54$ | $961.3 \pm 378.0$ |
| | Monocle3 | $0.995 \pm 0.002$ | $0.991 \pm 0.003$ | $7.00 \pm 0.00$ | $20.5 \pm 1.8$ |
| | SC3 | $0.000 \pm 0.000$ | $0.000 \pm 0.000$ | $1.00 \pm 0.00$ | $72.6 \pm 5.4$ |
| $K_{\text{true}} = 9$ | ProClust | $0.914 \pm 0.095$ | $0.913 \pm 0.093$ | $8.92 \pm 0.80$ | $506.3 \pm 65.1$ |
| | CHOIR | $0.512 \pm 0.348$ | $0.607 \pm 0.304$ | $8.34 \pm 4.89$ | $5880.3 \pm 2402.7$ |
| | scSHC | $0.615 \pm 0.322$ | $0.638 \pm 0.310$ | $11.28 \pm 5.18$ | $992.9 \pm 392.7$ |
| | Monocle3 | $0.938 \pm 0.040$ | $0.929 \pm 0.026$ | $8.86 \pm 0.38$ | $19.3 \pm 1.6$ |
| | SC3 | $0.000 \pm 0.000$ | $0.000 \pm 0.000$ | $1.00 \pm 0.00$ | $92.6 \pm 200.1$ |

Table S2: **Clustering performance in the PBMC-based semi-synthetic simulation.** The ground-truth number of clusters was $K_{\text{true}} = 3, 5, 7$, or 9. Values are mean ± standard deviation across 100 replicates per setting. Runtime is reported in seconds.

| Setting | Method | ARI | NMI | $\widehat{K}$ | Runtime (s) |
|---|---|---|---|---|---|
| Sim. 2: $K_{\text{true}} = 9$ | ProClust | $0.969 \pm 0.005$ | $0.972 \pm 0.003$ | $9.00 \pm 0.00$ | $532.5 \pm 33.2$ |
| | CHOIR | $0.875 \pm 0.035$ | $0.880 \pm 0.028$ | $11.14 \pm 0.84$ | $800.7 \pm 534.8$ |
| | scSHC | $0.907 \pm 0.005$ | $0.912 \pm 0.004$ | $10.01 \pm 0.10$ | $379.9 \pm 40.4$ |
| | Monocle3 | $0.723 \pm 0.025$ | $0.839 \pm 0.013$ | $8.01 \pm 0.10$ | $16.7 \pm 1.3$ |
| | SC3 | $0.532 \pm 0.057$ | $0.616 \pm 0.010$ | $43.44 \pm 0.76$ | $9266.7 \pm 620.6$ |
| Sim. 3: $K_{\text{true}} = 3$ | ProClust | $0.956 \pm 0.011$ | $0.936 \pm 0.016$ | $3.06 \pm 0.24$ | $241.9 \pm 24.6$ |
| | CHOIR | $0.819 \pm 0.079$ | $0.758 \pm 0.066$ | $4.46 \pm 0.67$ | $565.3 \pm 181.0$ |
| | scSHC | $0.761 \pm 0.094$ | $0.713 \pm 0.070$ | $4.70 \pm 0.64$ | $224.6 \pm 54.6$ |
| | Monocle3 | $0.534 \pm 0.000$ | $0.555 \pm 0.000$ | $2.00 \pm 0.00$ | $15.8 \pm 0.7$ |
| | SC3 | $0.421 \pm 0.090$ | $0.345 \pm 0.027$ | $47.11 \pm 0.67$ | $9740.5 \pm 506.4$ |
| Sim. 3: $K_{\text{true}} = 5$ | ProClust | $0.960 \pm 0.005$ | $0.958 \pm 0.005$ | $5.00 \pm 0.00$ | $354.7 \pm 15.8$ |
| | CHOIR | $0.857 \pm 0.054$ | $0.846 \pm 0.040$ | $6.69 \pm 0.72$ | $564.8 \pm 112.7$ |
| | scSHC | $0.890 \pm 0.010$ | $0.877 \pm 0.011$ | $6.04 \pm 0.20$ | $254.7 \pm 30.0$ |
| | Monocle3 | $0.689 \pm 0.001$ | $0.776 \pm 0.001$ | $4.00 \pm 0.00$ | $16.2 \pm 1.5$ |
| | SC3 | $0.470 \pm 0.064$ | $0.480 \pm 0.014$ | $45.44 \pm 0.64$ | $9760.8 \pm 832.9$ |
| Sim. 3: $K_{\text{true}} = 7$ | ProClust | $0.967 \pm 0.005$ | $0.969 \pm 0.004$ | $7.00 \pm 0.00$ | $397.8 \pm 30.2$ |
| | CHOIR | $0.873 \pm 0.046$ | $0.875 \pm 0.034$ | $9.20 \pm 0.95$ | $603.6 \pm 111.4$ |
| | scSHC | $0.899 \pm 0.054$ | $0.903 \pm 0.034$ | $7.98 \pm 0.32$ | $342.4 \pm 50.8$ |
| | Monocle3 | $0.718 \pm 0.003$ | $0.828 \pm 0.006$ | $5.99 \pm 0.10$ | $16.0 \pm 1.0$ |
| | SC3 | $0.523 \pm 0.054$ | $0.596 \pm 0.012$ | $43.23 \pm 0.57$ | $8926.3 \pm 558.6$ |

#### S3.2 Resolution sensitivity of Leiden and Louvain clustering

For the Monocle3 benchmark, we used its built-in Leiden clustering with `resolution = NULL`. We additionally examined the resolution sensitivity of Leiden and Louvain in both the parametric and PBMC-based semi-synthetic simulations for $K_{\text{true}} = 3, 5, 7, 9$. Resolutions ranged from 0.1 to 2.0 in increments of 0.1, with 100 independent simulation replicates per setting. Raw counts were log-normalized with a scaling factor of $10^4$, followed by selection of 2,000 highly variable features, standardization, and PCA. Both methods used an SNN graph constructed from the first 30 principal components with $k = 10$. Mean ARI and mean estimated cluster counts are shown in Supplementary Figures S1 and S2.

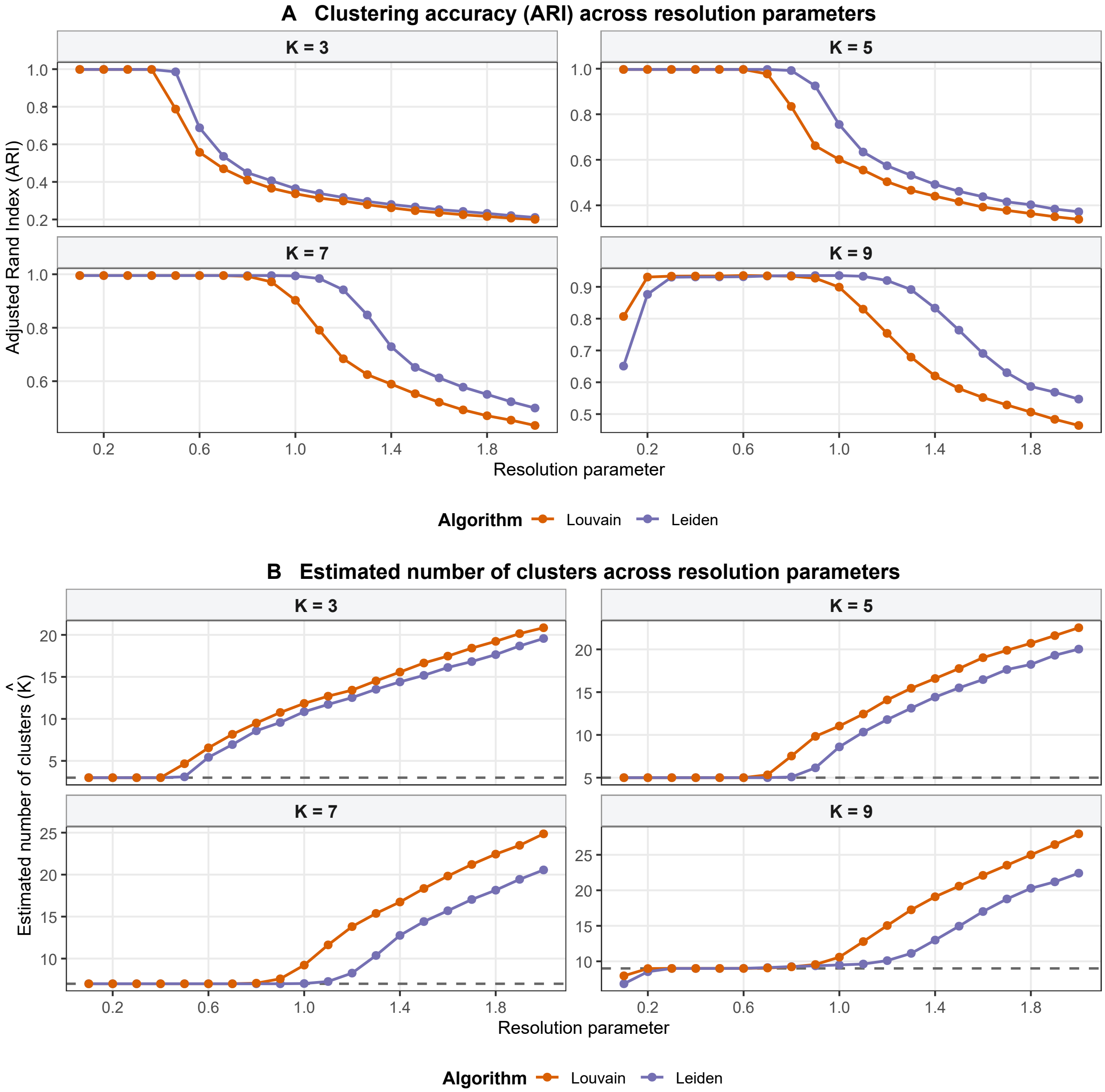


Figure S1: **Resolution sensitivity of Leiden and Louvain in the parametric simulation. A**, Mean adjusted Rand index (ARI) across clustering resolutions from 0.1 to 2.0 in increments of 0.1 for $K_{\text{true}} = 3, 5, 7, 9$. **B**, Mean estimated number of clusters ($\widehat{K}$) for the same settings. Each point represents the mean across 100 independent simulation replicates. Dashed horizontal lines indicate the ground-truth number of clusters. Orange and purple curves represent Louvain and Leiden, respectively.

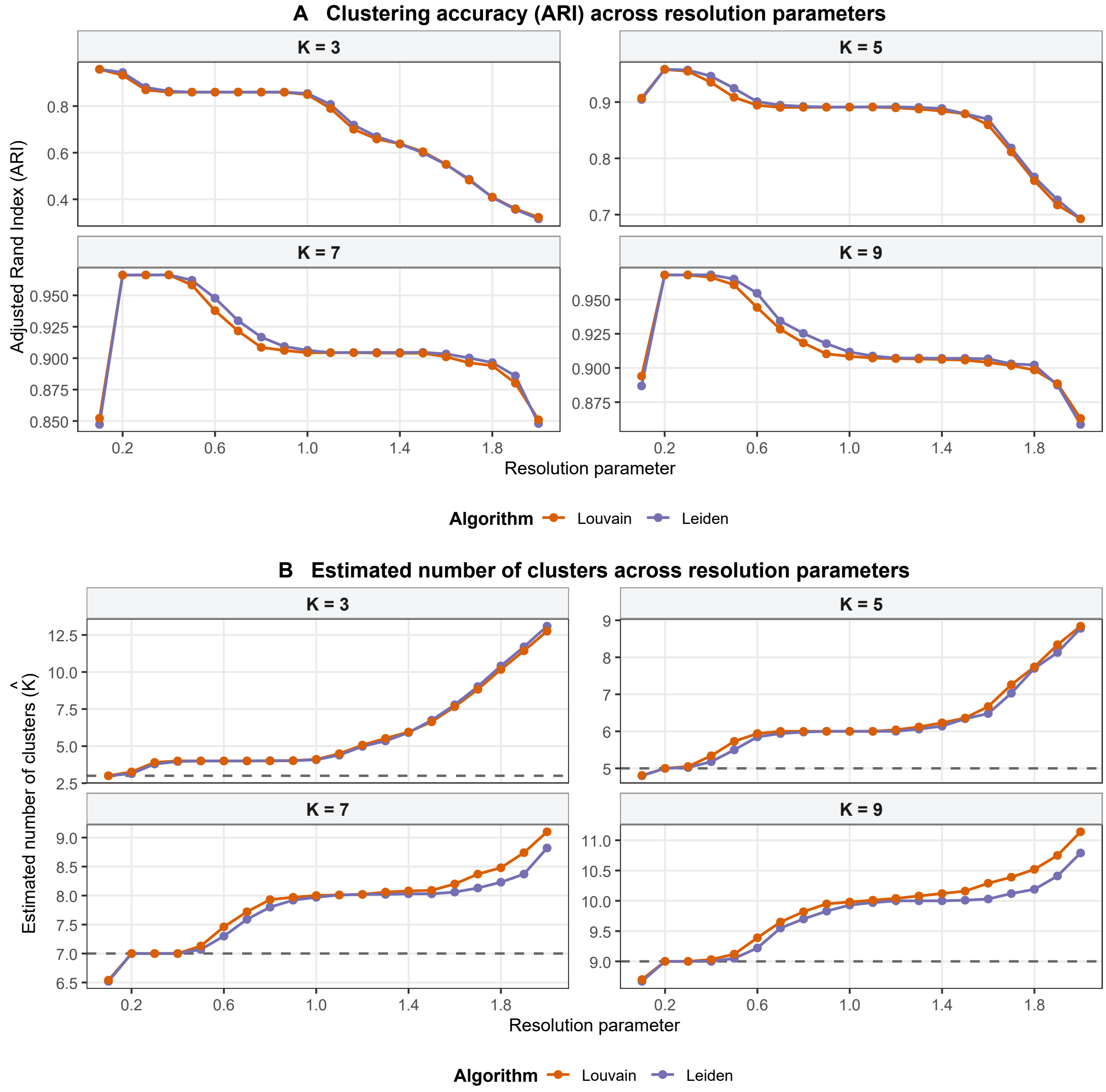


Figure S2: **Resolution sensitivity of Leiden and Louvain in the PBMC-based semi-synthetic simulation.** **A**, Mean adjusted Rand index (ARI) across clustering resolutions from 0.1 to 2.0 in increments of 0.1 for $K_{\text{true}} = 3, 5, 7, 9$. **B**, Mean estimated number of clusters $(\widehat{K})$ for the same settings. Each point represents the mean across 100 independent simulation replicates. Dashed horizontal lines indicate the ground-truth number of clusters. Orange and purple curves represent Louvain and Leiden, respectively.

### S4 Additional Tabula Muris Results

#### S4.1 Brain non-myeloid

The brain non-myeloid dataset contained 3,401 cells representing seven reference-annotated cell types. Clustering performance metrics, estimated cluster counts, and runtimes are reported in Supplementary Table S3.

Table S3: **Clustering performance on the Tabula Muris brain non-myeloid dataset.** The reference annotation contained seven cell types. ProClust was initialized from an over-clustered partition of $K_{\text{initial}} = 20$ clusters. Runtime is reported in seconds.

| Method | ARI | NMI | $\widehat{K}$ | Runtime (s) |
|---|---|---|---|---|
| ProClust | 0.824 | 0.766 | 10 | 369.87 |
| Leiden | 0.470 | 0.607 | 14 | 0.47 |
| Louvain | 0.458 | 0.591 | 17 | 0.19 |
| CHOIR | 0.533 | 0.579 | 18 | 278.60 |
| scSHC | 0.727 | 0.675 | 12 | 364.44 |
| Monocle3 | 0.820 | 0.800 | 10 | 15.92 |
| SC3 | 0.287 | 0.469 | 26 | 1036.55 |

ProClust identified 10 final clusters and achieved an ARI of 0.824 and an NMI of 0.766. Leiden and Louvain, both run at resolution 0.8, identified 14 and 17 clusters, with ARIs of 0.470 and 0.458, respectively. CHOIR identified 18 clusters (ARI = 0.533), scSHC identified 12 clusters (ARI = 0.727), and SC3 identified 26 clusters (ARI = 0.287). Monocle3 also identified 10 clusters, achieving an ARI of 0.820 and an NMI of 0.800. ProClust achieved the highest ARI, whereas Monocle3 achieved the highest NMI among the evaluated methods.

The initial clustering partition, final ProClust clusters, and reference cell-type annotations are visualized on the same UMAP embedding in Supplementary Figure S3.

The hierarchical testing process is summarized in Supplementary Figure S4. ProClust reduced the initial partition of 20 clusters to 10 final clusters through sequential node-level hypothesis testing. The alignment between the final ProClust clusters and the reference cell-type annotations is shown in Supplementary Figure S5.

The cluster-to-reference composition heatmap showed that clusters corresponding to oligodendrocyte precursor cells, brain pericytes, and astrocytes were predominantly composed of cells with the matching reference annotations. Reference-annotated neurons, oligodendrocytes, and endothelial cells were distributed across multiple ProClust clusters, indicating a finer partition than the reference annotation. The transcriptional differences between the two endothelial clusters are characterized

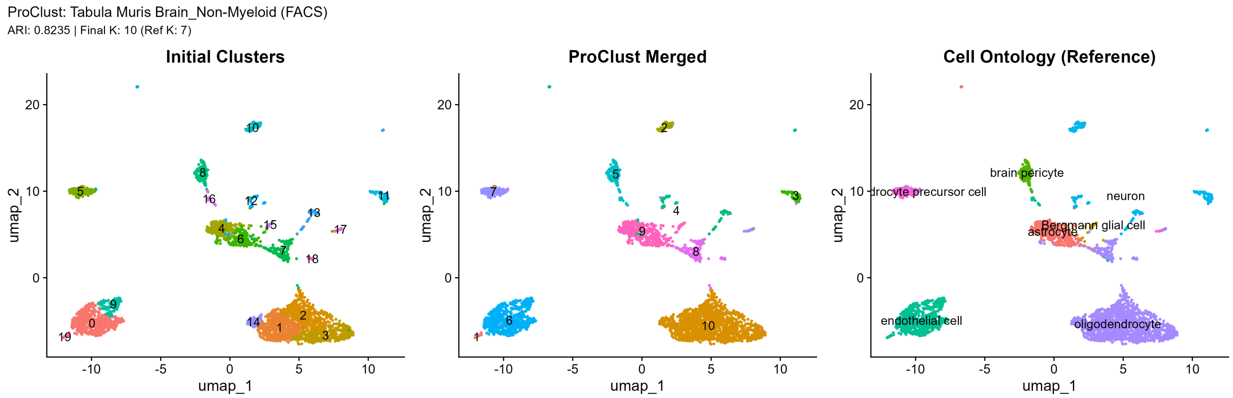


Figure S3: **UMAP visualization of clustering results for the Tabula Muris brain non-myeloid dataset.** The same UMAP embedding is colored by the initial clustering partition (left), the final ProClust clusters (middle), and the reference cell-type annotations (right). Numeric labels in the left and middle panels indicate initial and final cluster identifiers, respectively.

117 in the main text.

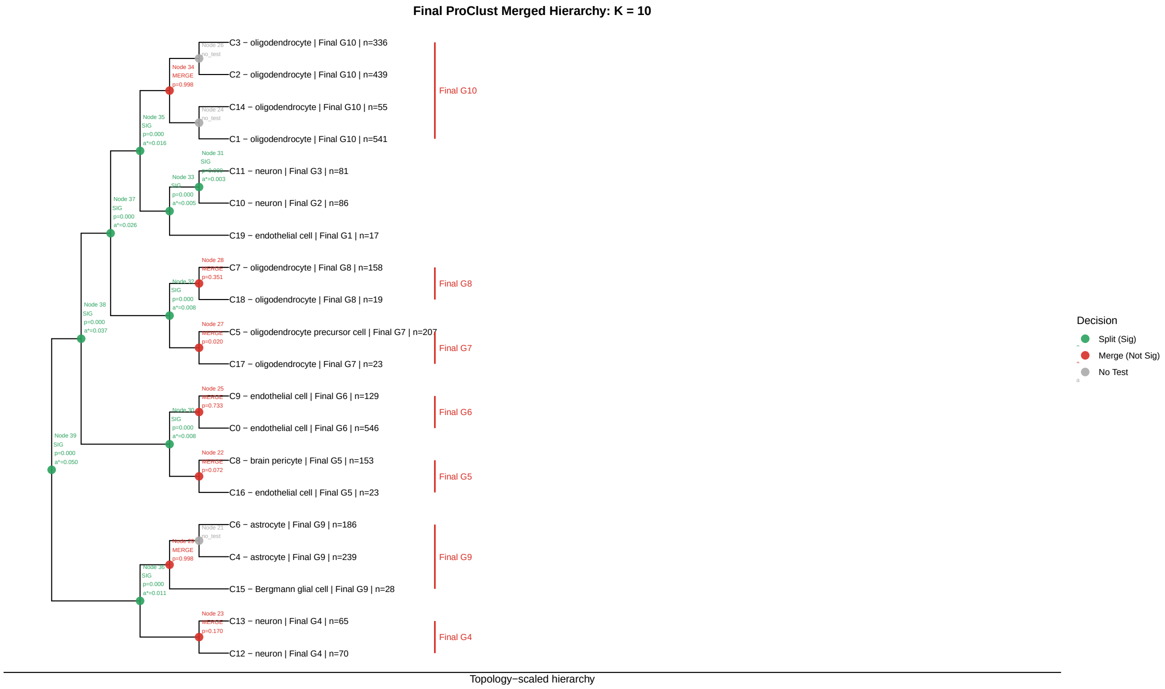


Figure S4: **Hierarchical testing results for the Tabula Muris brain non-myeloid dataset.** ProClust reduced an initial partition of 20 clusters to 10 final clusters through sequential node-level hypothesis testing. Green nodes indicate statistically supported splits, red nodes indicate nonsignificant splits for which the corresponding child branches were merged, and gray nodes indicate nodes at which no test was performed. For each tested node, the node identifier, $p$-value, and node-specific threshold $\alpha^*$ are shown. Leaf labels report the initial cluster identifier, dominant reference cell-type annotation, final ProClust cluster, and number of cells.

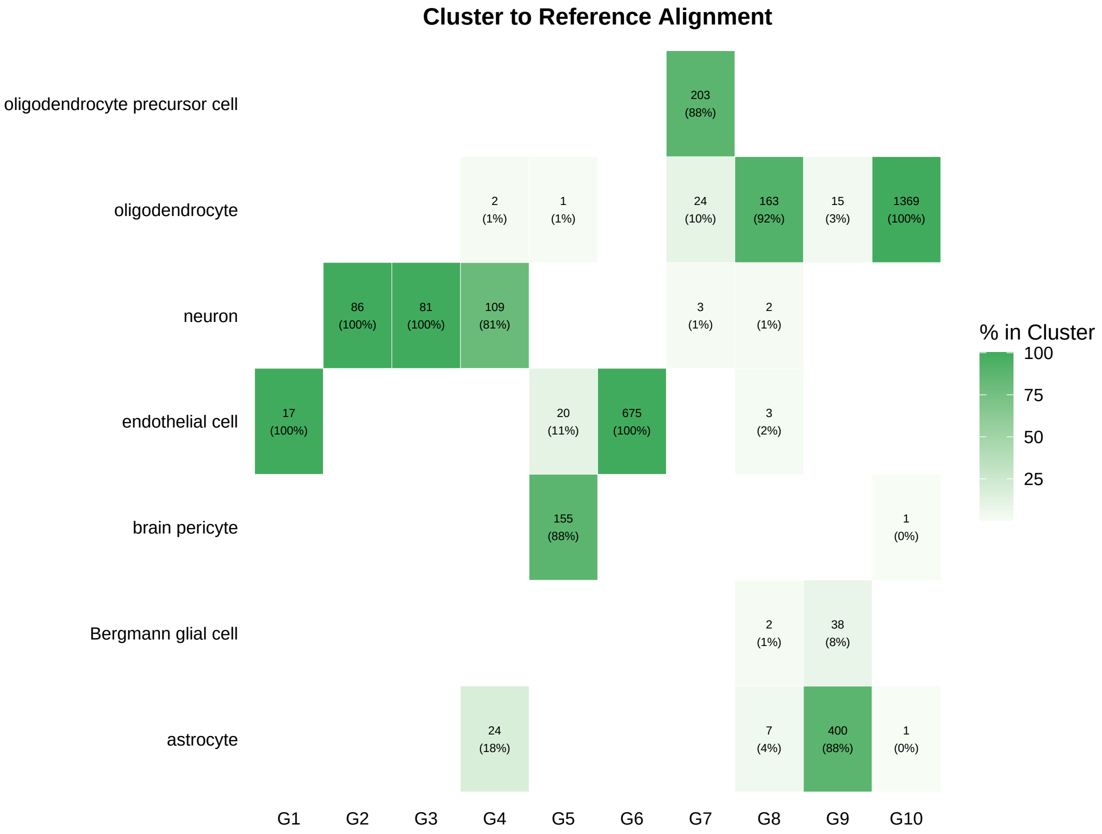


Figure S5: **Alignment between the final ProClust clusters and reference cell-type annotations in the Tabula Muris brain non-myeloid dataset.** Columns correspond to the 10 final ProClust clusters, and rows correspond to the reference cell-type annotations. Each nonempty cell reports the number of cells and the percentage of the corresponding ProClust cluster assigned to that reference annotation. Darker shading indicates a larger percentage within the ProClust cluster.

#### S4.2 Lung

The lung dataset contained 1,676 cells representing 11 reference-annotated cell types. Clustering performance metrics, estimated cluster counts, and runtimes are reported in Supplementary Table S4.

Table S4: **Clustering performance on the Tabula Muris lung dataset.** The reference annotation contained 11 cell types. ProClust was initialized from an over-clustered partition of $K_{\text{initial}} = 20$ clusters. Runtime is reported in seconds.

| Method | ARI | NMI | $\widehat{K}$ | Runtime (s) |
|---|---|---|---|---|
| ProClust | 0.534 | 0.673 | 15 | 312.25 |
| Leiden | 0.368 | 0.606 | 16 | 0.12 |
| Louvain | 0.497 | 0.656 | 18 | 0.06 |
| CHOIR | 0.335 | 0.583 | 21 | 186.94 |
| scSHC | 0.379 | 0.616 | 17 | 456.28 |
| Monocle3 | 0.609 | 0.699 | 14 | 13.14 |
| SC3 | 0.292 | 0.558 | 24 | 2334.28 |

Starting from an initial partition of 20 clusters, ProClust achieved an ARI of 0.534 and NMI of 0.673, estimating $\widehat{K} = 15$ final clusters. In comparison, graph-based methods such as Leiden and Louvain at resolution 0.8 yielded ARIs of 0.368 ($\widehat{K} = 16$) and 0.497 ($\widehat{K} = 18$), respectively. CHOIR identified 21 clusters (ARI = 0.335), scSHC estimated 17 clusters (ARI = 0.379), and SC3 produced 24 clusters (ARI = 0.292). Monocle3 attained an ARI of 0.609 ($\widehat{K} = 14$).

The initial clustering partition, final ProClust clusters, and reference cell-type annotations are visualized on the same UMAP embedding in Supplementary Figure S6.

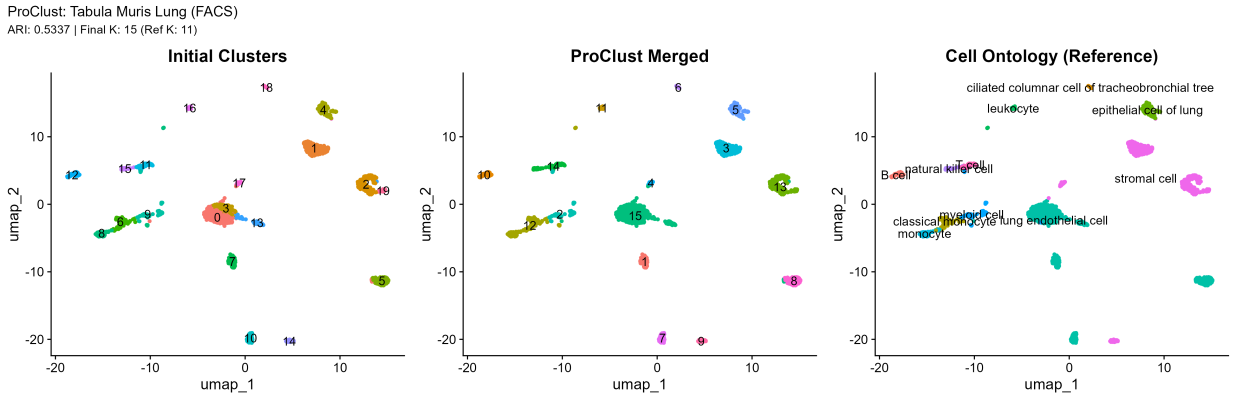


Figure S6: **UMAP visualization of clustering results for the Tabula Muris lung dataset.** The same UMAP embedding is colored by the initial clustering partition (left), the final ProClust clusters (middle), and the reference cell-type annotations (right). Numeric labels in the left and middle panels indicate initial and final cluster identifiers, respectively.

The hierarchical testing process for the lung dataset is summarized in Supplementary Figure S7. ProClust reduced the initial partition of 20 clusters to 15 final clusters through sequential node-level hypothesis testing. The alignment between the final ProClust clusters and the reference cell-type annotations is shown in Supplementary Figure S8.

The cluster-to-reference composition heatmap showed that clusters corresponding to lung epithelial cells, ciliated columnar cells of the tracheobronchial tree, and B cells were predominantly composed of cells with the matching reference annotations. Reference-annotated lung endothelial and stromal cells were distributed across multiple ProClust clusters, indicating a finer partition within these broad cell categories. Node-level testing and marker-gene analyses supporting the retained endothelial and stromal subdivisions are presented in the main text.

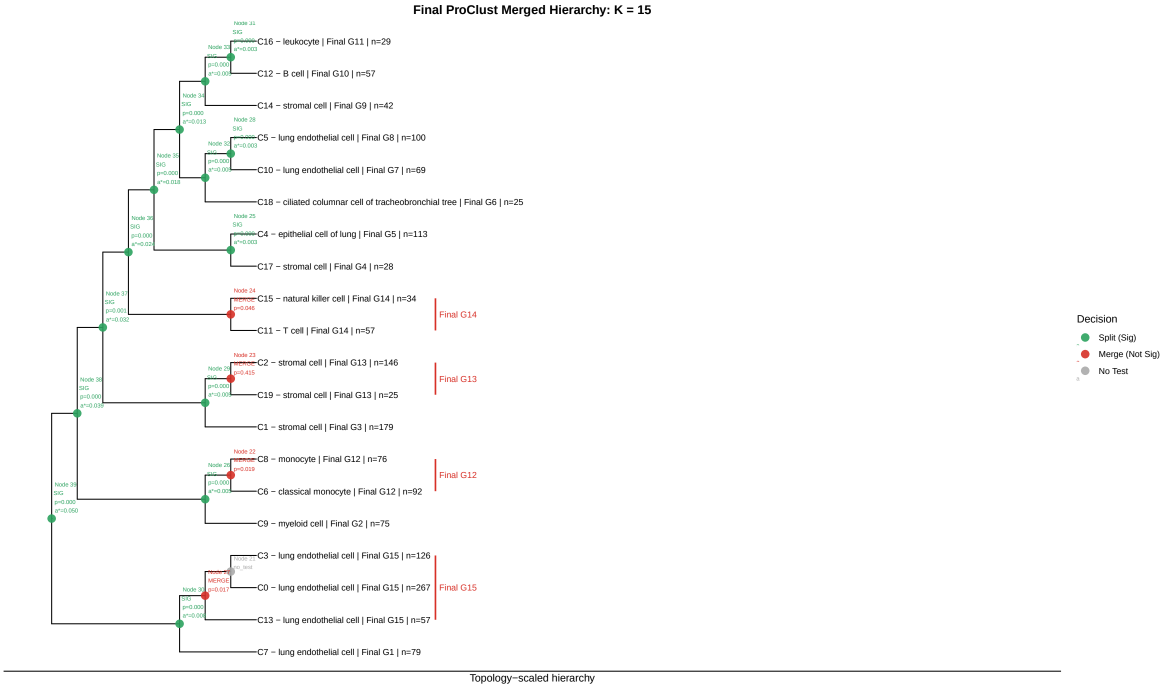


Figure S7: **Hierarchical testing results for the Tabula Muris lung dataset.** ProClust reduced an initial partition of 20 clusters to 15 final clusters through sequential node-level hypothesis testing. Green nodes indicate statistically supported splits, red nodes indicate nonsignificant splits for which the corresponding child branches were merged, and gray nodes indicate nodes at which no test was performed. For each tested node, the node identifier, $p$-value, and node-specific threshold $\alpha^*$ are shown. Leaf labels report the initial cluster identifier, dominant reference cell-type annotation, final ProClust cluster, and number of cells.

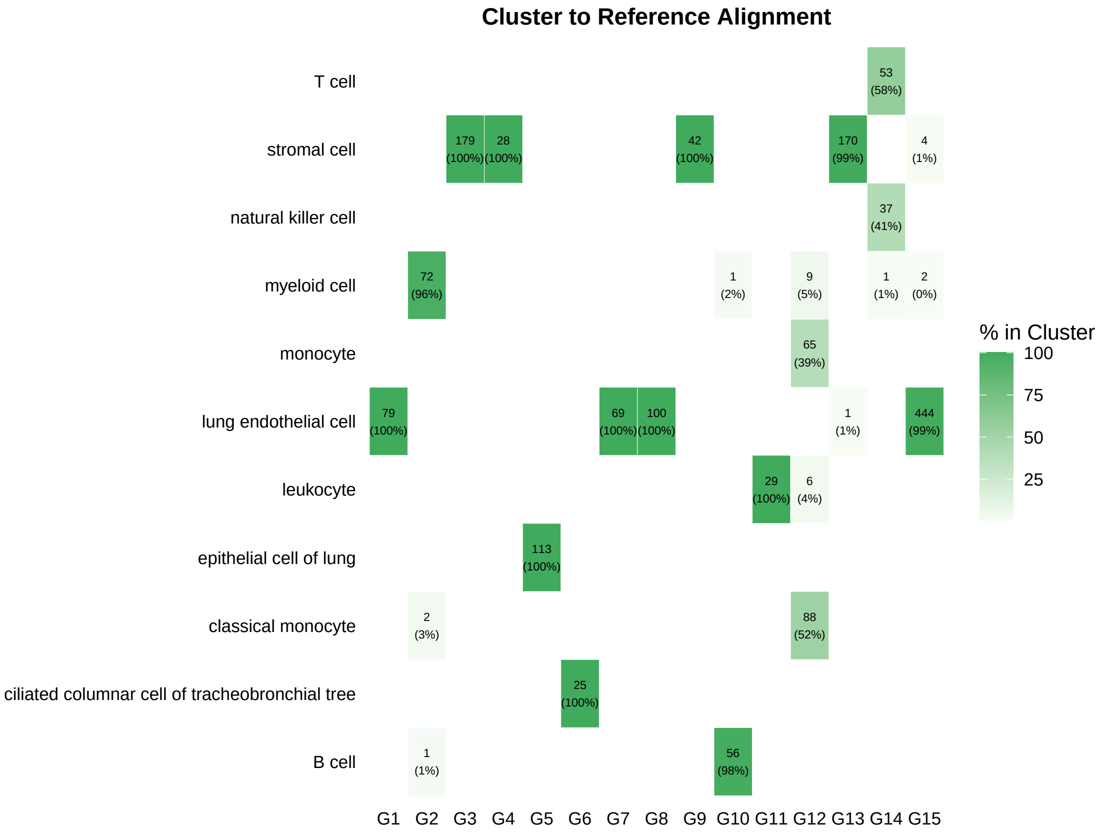


Figure S8: **Alignment between the final ProClust clusters and reference cell-type annotations in the Tabula Muris lung dataset.** Columns correspond to the 15 final ProClust clusters, and rows correspond to the reference cell-type annotations. Each nonempty cell reports the number of cells and the percentage of the corresponding ProClust cluster assigned to that reference annotation. Darker shading indicates a larger percentage within the ProClust cluster.

#### S4.3 Pancreas

The pancreas dataset contained 1,564 cells representing nine reference-annotated cell types. Clustering performance metrics, estimated cluster counts, and runtimes are reported in Supplementary Table S5.

Table S5: **Clustering performance on the Tabula Muris pancreas dataset.** The reference annotation contained nine cell types. ProClust was initialized from an over-clustered partition of $K_{\text{initial}} = 15$ clusters. Runtime is reported in seconds.

| Method | ARI | NMI | $\widehat{K}$ | Runtime (s) |
|---|---|---|---|---|
| ProClust | 0.645 | 0.746 | 12 | 166.80 |
| Leiden | 0.647 | 0.749 | 11 | 0.24 |
| Louvain | 0.618 | 0.712 | 14 | 0.11 |
| CHOIR | 0.442 | 0.629 | 22 | 206.14 |
| scSHC | 0.894 | 0.890 | 10 | 350.97 |
| Monocle3 | 0.534 | 0.682 | 13 | 13.09 |
| SC3 | 0.704 | 0.763 | 17 | 1315.94 |

Starting from an initial partition of 15 clusters, ProClust achieved an ARI of 0.645 and NMI of 0.746, estimating $\widehat{K} = 12$ final clusters. In comparison, graph-based methods such as Leiden and Louvain at resolution 0.8 yielded ARIs of 0.647 ($\widehat{K} = 11$) and 0.618 ($\widehat{K} = 14$), respectively. CHOIR identified 22 clusters (ARI = 0.442), scSHC estimated 10 clusters (ARI = 0.894), and SC3 produced 17 clusters (ARI = 0.704). Monocle3 attained an ARI of 0.534 ($\widehat{K} = 13$).

The initial clustering partition, final ProClust clusters, and reference cell-type annotations are visualized on the same UMAP embedding in Supplementary Figure S9.

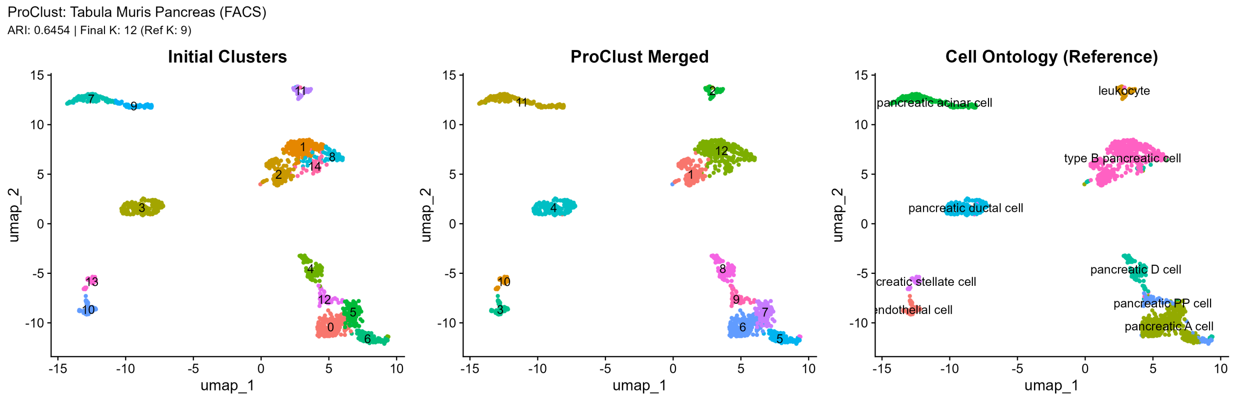


Figure S9: **UMAP visualization of clustering results for the Tabula Muris pancreas dataset.** The same UMAP embedding is colored by the initial clustering partition (left), the final ProClust clusters (middle), and the reference cell-type annotations (right). Numeric labels in the left and middle panels indicate initial and final cluster identifiers, respectively.

The hierarchical testing process for the pancreas dataset is summarized in Supplementary Figure S10. ProClust reduced the initial partition of 15 clusters to 12 final clusters through sequential node-level hypothesis testing. The alignment between the final ProClust clusters and the reference cell-type annotations is shown in Supplementary Figure S11.

The cluster-to-reference composition heatmap summarizes the reference cell-type composition of each of the 12 final ProClust clusters. ProClust identified 12 final clusters, compared with nine reference-annotated cell types. Their correspondence with the reference annotations is shown in Supplementary Figure S11.

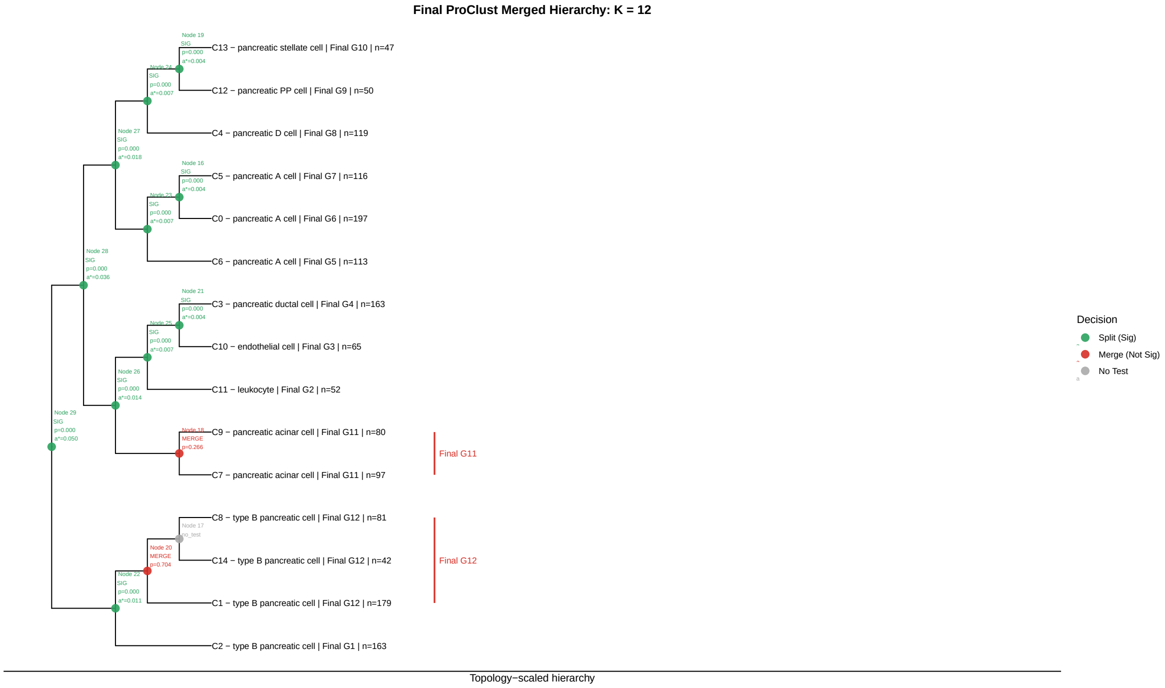


Figure S10: **Hierarchical testing results for the Tabula Muris pancreas dataset.** ProClust reduced an initial partition of 15 clusters to 12 final clusters through sequential node-level hypothesis testing. Green nodes indicate statistically supported splits, red nodes indicate nonsignificant splits for which the corresponding child branches were merged, and gray nodes indicate nodes at which no test was performed. For each tested node, the node identifier, $p$-value, and node-specific threshold $\alpha^*$ are shown. Leaf labels report the initial cluster identifier, dominant reference cell-type annotation, final ProClust cluster, and number of cells.

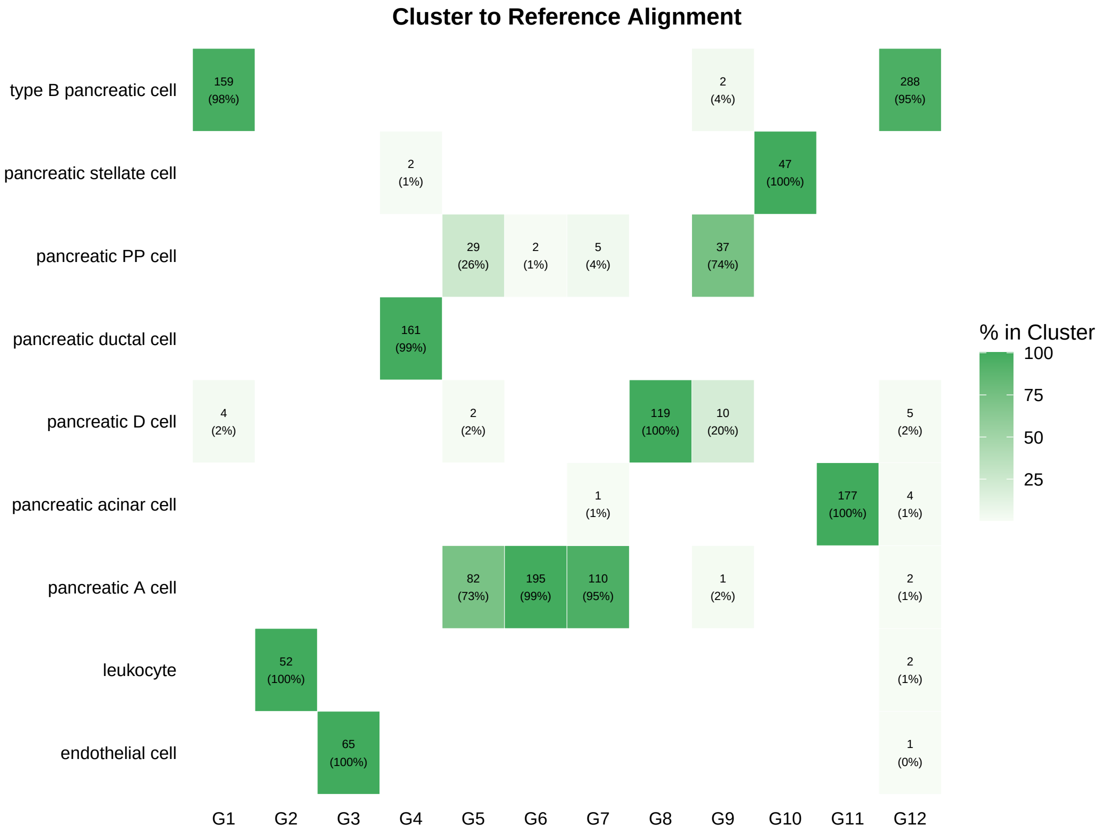


Figure S11: **Alignment between the final ProClust clusters and reference cell-type annotations in the Tabula Muris pancreas dataset.** Columns correspond to the 12 final ProClust clusters, and rows correspond to the reference cell-type annotations. Each nonempty cell reports the number of cells and the percentage of the corresponding ProClust cluster assigned to that reference annotation. Darker shading indicates a larger percentage within the ProClust cluster.

#### S4.4 Spleen

The spleen dataset contained 1,697 cells representing three reference-annotated cell types. Clustering performance metrics, estimated cluster counts, and runtimes are reported in Supplementary Table S6.

Table S6: **Clustering performance on the Tabula Muris spleen dataset.** The reference annotation contained three cell types. ProClust was initialized from an over-clustered partition of $K_{\text{initial}} = 11$ clusters. Runtime is reported in seconds.

| Method | ARI | NMI | $\widehat{K}$ | Runtime (s) |
|---|---|---|---|---|
| ProClust | 0.884 | 0.795 | 3 | 109.70 |
| Leiden | 0.262 | 0.357 | 7 | 0.21 |
| Louvain | 0.283 | 0.352 | 8 | 0.11 |
| CHOIR | 0.333 | 0.423 | 5 | 123.61 |
| scSHC | 0.000 | 0.000 | 1 | 152.63 |
| Monocle3 | 0.615 | 0.549 | 3 | 9.44 |
| SC3 | 0.106 | 0.238 | 20 | 3843.55 |

ProClust identified three final clusters and achieved an ARI of 0.884 and an NMI of 0.795. Leiden and Louvain, both run at resolution 0.8, identified seven and eight clusters, with ARIs of 0.262 and 0.283, respectively. CHOIR identified five clusters (ARI = 0.333), and SC3 identified 20 clusters (ARI = 0.106). Under the evaluated settings, scSHC returned a single cluster, with both ARI and NMI equal to zero. Monocle3 identified three clusters, with an ARI of 0.615.

The initial clustering partition, final ProClust clusters, and reference cell-type annotations are visualized on the same UMAP embedding in Supplementary Figure S12.

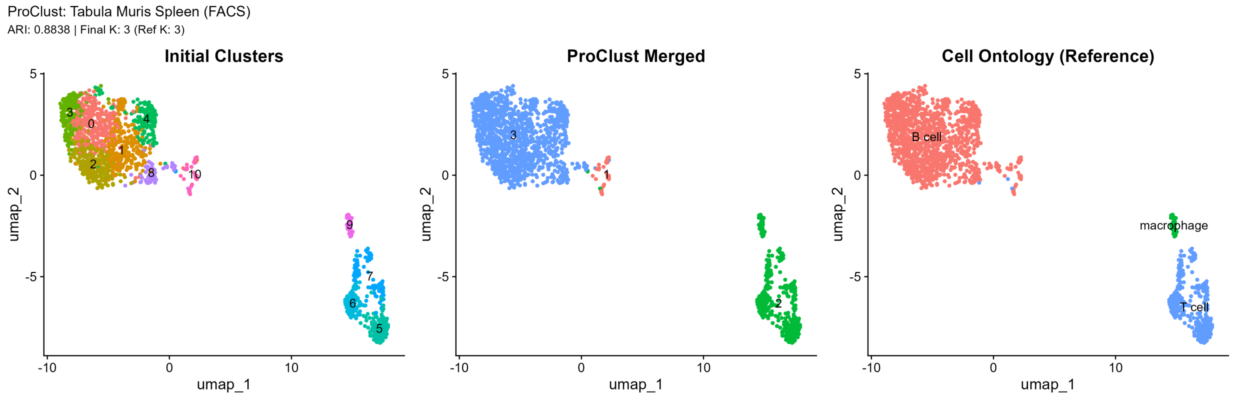


Figure S12: **UMAP visualization of clustering results for the Tabula Muris spleen dataset.** The same UMAP embedding is colored by the initial clustering partition (left), the final ProClust clusters (middle), and the reference cell-type annotations (right). Numeric labels in the left and middle panels indicate initial and final cluster identifiers, respectively.

The hierarchical testing process for the spleen dataset is summarized in Supplementary Figure S13. ProClust reduced the initial partition of 11 clusters to three final clusters through sequential node-level hypothesis testing. The alignment between the final ProClust clusters and the reference cell-type annotations is shown in Supplementary Figure S14.

ProClust retained a large B-cell-dominated cluster (cluster 3) and a smaller B-cell-dominated cluster (cluster 1), while reference-annotated T cells and macrophages were grouped together in cluster 2. Although ProClust identified three final clusters, matching the number of reference-annotated cell types, the resulting partition did not correspond one-to-one with the reference annotation.

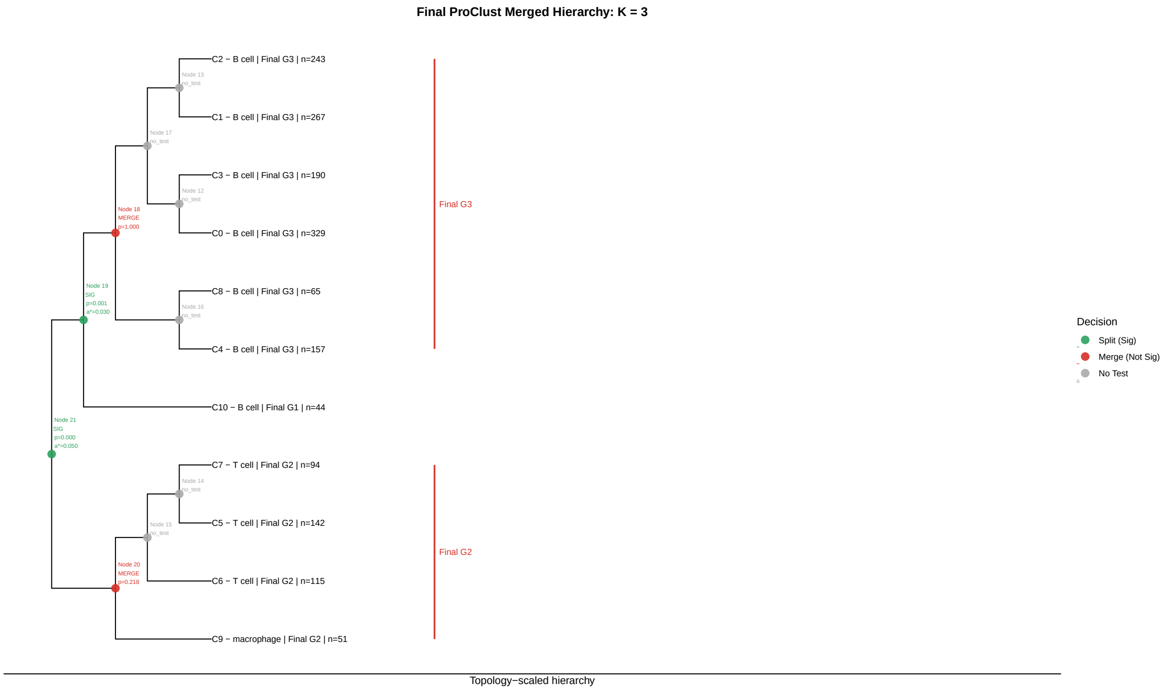


Figure S13: **Hierarchical testing results for the Tabula Muris spleen dataset.** ProClust reduced an initial partition of 11 clusters to 3 final clusters through sequential node-level hypothesis testing. Green nodes indicate statistically supported splits, red nodes indicate nonsignificant splits for which the corresponding child branches were merged, and gray nodes indicate nodes at which no test was performed. For each tested node, the node identifier, $p$-value, and node-specific threshold $\alpha^*$ are shown. Leaf labels report the initial cluster identifier, dominant reference cell-type annotation, final ProClust cluster, and number of cells.

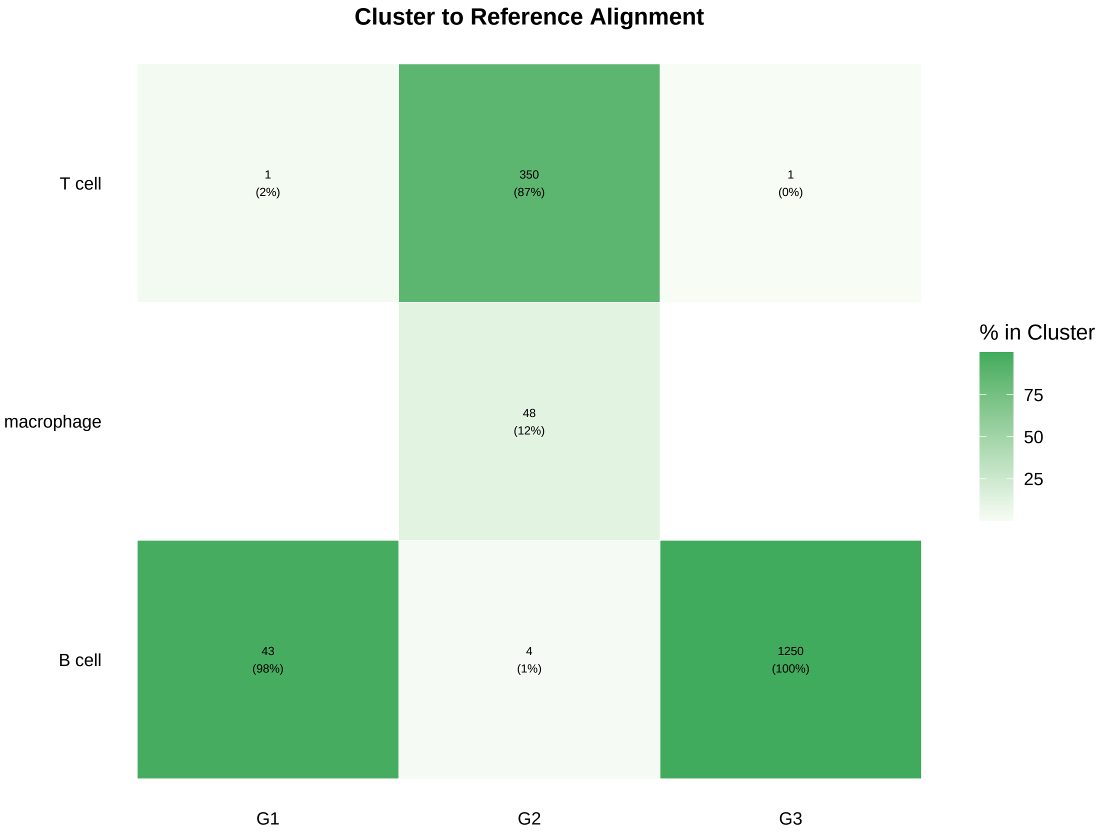


Figure S14: **Alignment between the final ProClust clusters and reference cell-type annotations in the Tabula Muris spleen dataset.** Columns correspond to the 3 final ProClust clusters, and rows correspond to the reference cell-type annotations. Each nonempty cell reports the number of cells and the percentage of the corresponding ProClust cluster assigned to that reference annotation. Darker shading indicates a larger percentage within the ProClust cluster.

#### S4.5 Heart

The heart dataset contained 4,337 cells representing seven reference-annotated cell types. Clustering performance metrics, estimated cluster counts, and runtimes are reported in Supplementary Table S7.

Table S7: **Clustering performance on the Tabula Muris heart dataset.** The reference annotation contained seven cell types. ProClust was initialized from an over-clustered partition of $K_{\text{initial}} = 21$ clusters. Runtime is reported in seconds.

| Method | ARI | NMI | $\widehat{K}$ | Runtime (s) |
|---|---|---|---|---|
| ProClust | 0.503 | 0.599 | 13 | 915.54 |
| Leiden | 0.409 | 0.554 | 14 | 0.69 |
| Louvain | 0.392 | 0.543 | 16 | 0.25 |
| CHOIR | 0.412 | 0.530 | 18 | 921.11 |
| scSHC | 0.343 | 0.496 | 14 | 820.59 |
| Monocle3 | 0.936 | 0.872 | 8 | 25.55 |
| SC3 | 0.212 | 0.408 | 41 | 3431.59 |

ProClust identified 13 final clusters and achieved an ARI of 0.503 and an NMI of 0.599. Leiden and Louvain, both run at resolution 0.8, identified 14 and 16 clusters, with ARIs of 0.409 and 0.392, respectively. CHOIR identified 18 clusters (ARI = 0.412), scSHC identified 14 clusters (ARI = 0.343), and SC3 identified 41 clusters (ARI = 0.212). Monocle3 identified eight clusters, with an ARI of 0.936.

The initial clustering partition, final ProClust clusters, and reference cell-type annotations are visualized on the same UMAP embedding in Supplementary Figure S15.

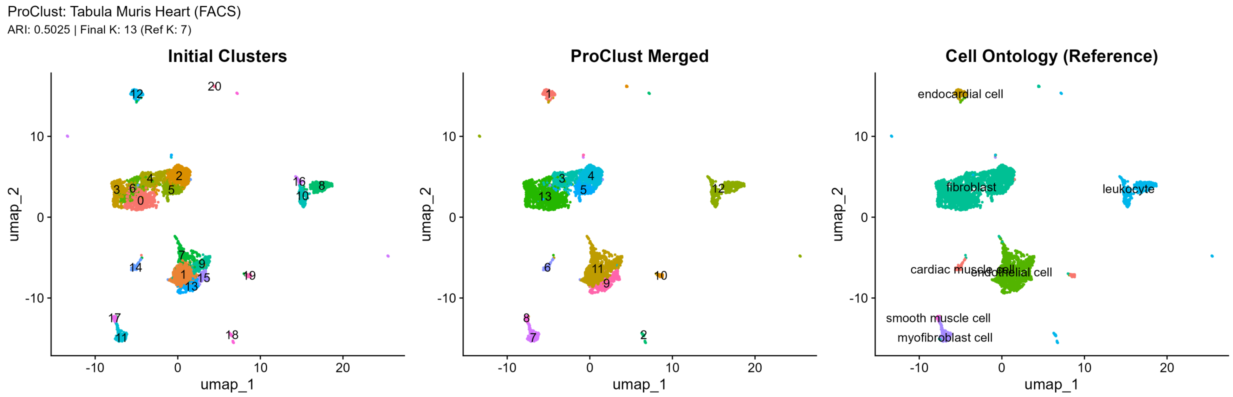


Figure S15: **UMAP visualization of clustering results for the Tabula Muris heart dataset.** The same UMAP embedding is colored by the initial clustering partition (left), the final ProClust clusters (middle), and the reference cell-type annotations (right). Numeric labels in the left and middle panels indicate initial and final cluster identifiers, respectively.

The hierarchical testing process for the heart dataset is summarized in Supplementary Figure S16. ProClust reduced the initial partition of 21 clusters to 13 final clusters through sequential node-level hypothesis testing. The alignment between the final ProClust clusters and the reference cell-type annotations is shown in Supplementary Figure S20.

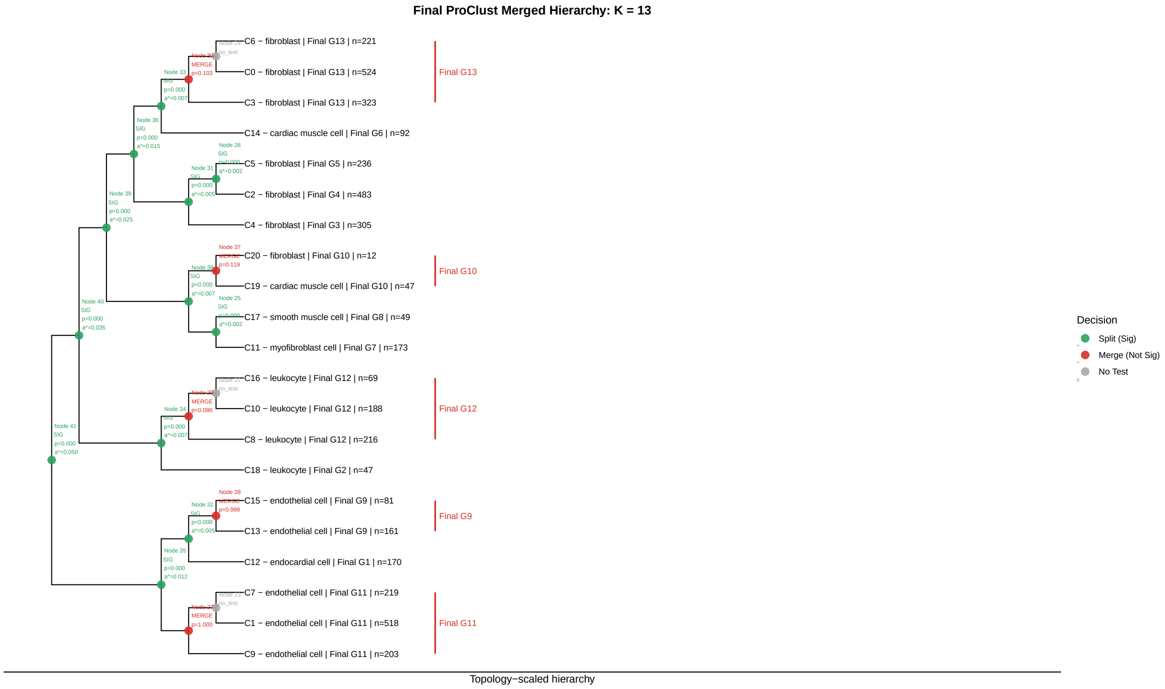


Figure S16: **Hierarchical testing results for the Tabula Muris heart dataset.** ProClust reduced an initial partition of 21 clusters to 13 final clusters through sequential node-level hypothesis testing. Green nodes indicate statistically supported splits, red nodes indicate nonsignificant splits for which the corresponding child branches were merged, and gray nodes indicate nodes at which no test was performed. For each tested node, the node identifier, $p$-value, and node-specific threshold $\alpha^*$ are shown. Leaf labels report the initial cluster identifier, dominant reference cell-type annotation, final ProClust cluster, and number of cells.

The heart dataset illustrates a limitation imposed by the fixed initial tree topology. Node 35 contained 1,352 cells, including initial cluster C12 (170 cells), annotated as endocardial cell, and clusters C15, C13, C7, C1, and C9, predominantly annotated as endothelial cells. In the UMAP embedding, C12 occupied a spatially distinct region, whereas the endothelial-associated clusters occupied a common region (Supplementary Figure S15). The three-group LFDA visualization at node 35 similarly showed one clearly separated group and substantial overlap between the other two (Supplementary Figure S17).

The initial hierarchy nevertheless placed $\{C12, C13, C15\}$ and $\{C1, C7, C9\}$ in two separate child branches (Supplementary Figure S16). The significant test result at node 35 provided evidence of clustering structure within the node, but the procedure continued along these predefined branches rather than using the test result to redefine the partition. Subsequent testing at node 32, which contained 412 cells, separated C12 from $\{C13, C15\}$, preserving a distinction consistent with the reference annotations (Supplementary Figure S18). In the other child branch, the test at node 27, which contained 940 cells, was nonsignificant, and $\{C1, C7, C9\}$ were merged into a single final cluster (Supplementary Figure S19). However, $\{C13, C15\}$ and $\{C1, C7, C9\}$ remained as two separate final clusters. Because merging is restricted to complete subtrees, the procedure could not merge these two endothelial-associated groups while retaining C12 as a separate cluster. This example shows that local tests can identify clustering structure within a subtree, while the final partition remains constrained by the initial tree topology. Such constraints may contribute to additional subdivision within a reference cell type, although a shared annotation and visual proximity alone do not establish that the groups should be merged.

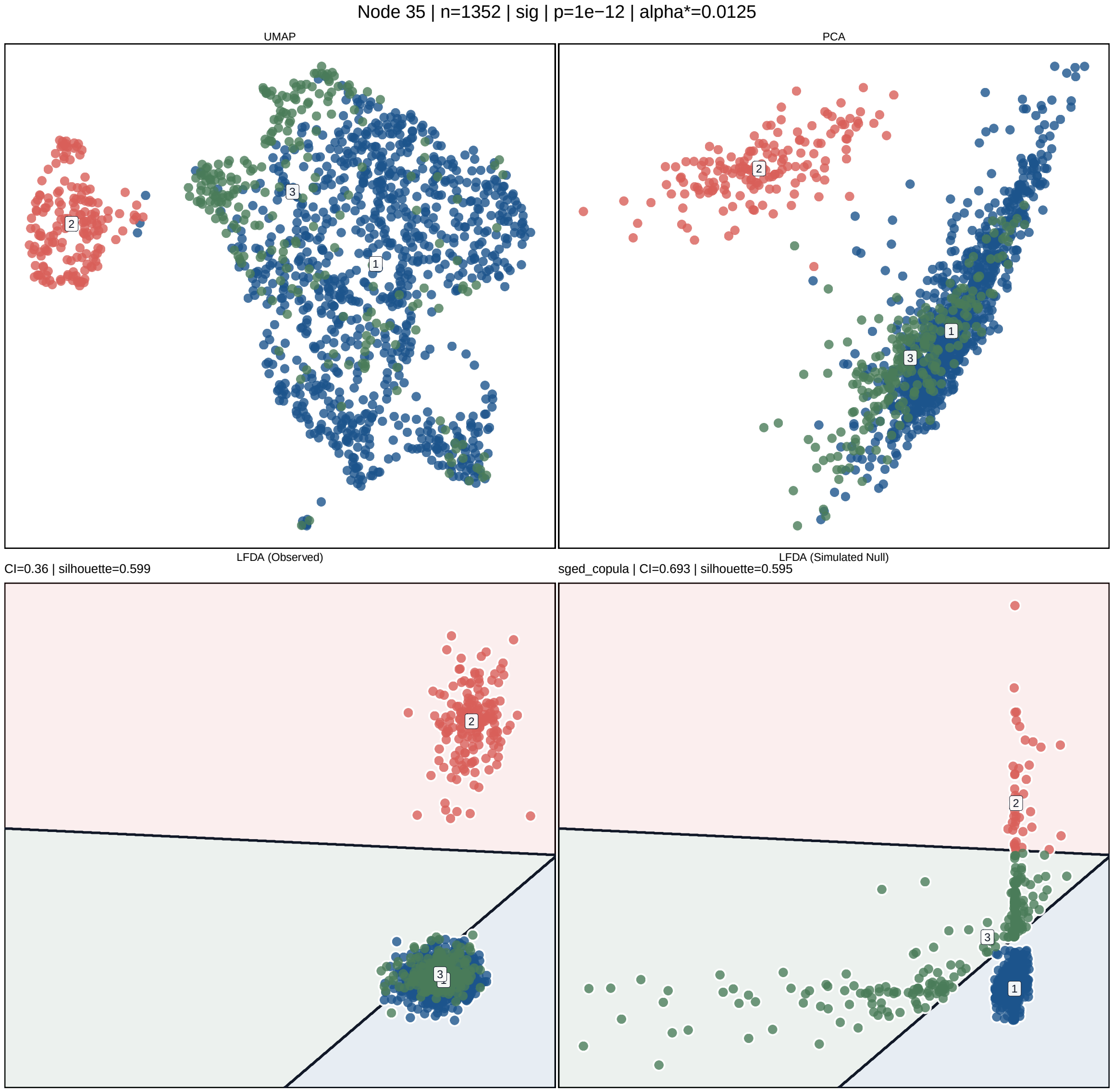


Figure S17: **Diagnostic visualization of node 35 in the Tabula Muris heart hierarchy.** Node 35 contained 1,352 cells. Panels show the UMAP embedding (top left), PCA space (top right), observed two-dimensional LFDA projection (bottom left), and a simulated sample from the fitted null model (bottom right). Colors and numeric labels identify the three groups used for the observed-data visualization; simulated cells are colored by their assigned decision regions. Shaded backgrounds and black lines in the LFDA panels indicate nearest-centroid decision regions and their boundaries. The observed LFDA projection shows one clearly separated group and substantial overlap between the other two. The node-level test was significant ($p = 10^{-12}$; $\alpha^* = 0.0125$), and testing continued along the predefined child branches. The displayed CI and silhouette values summarize the plotted samples.

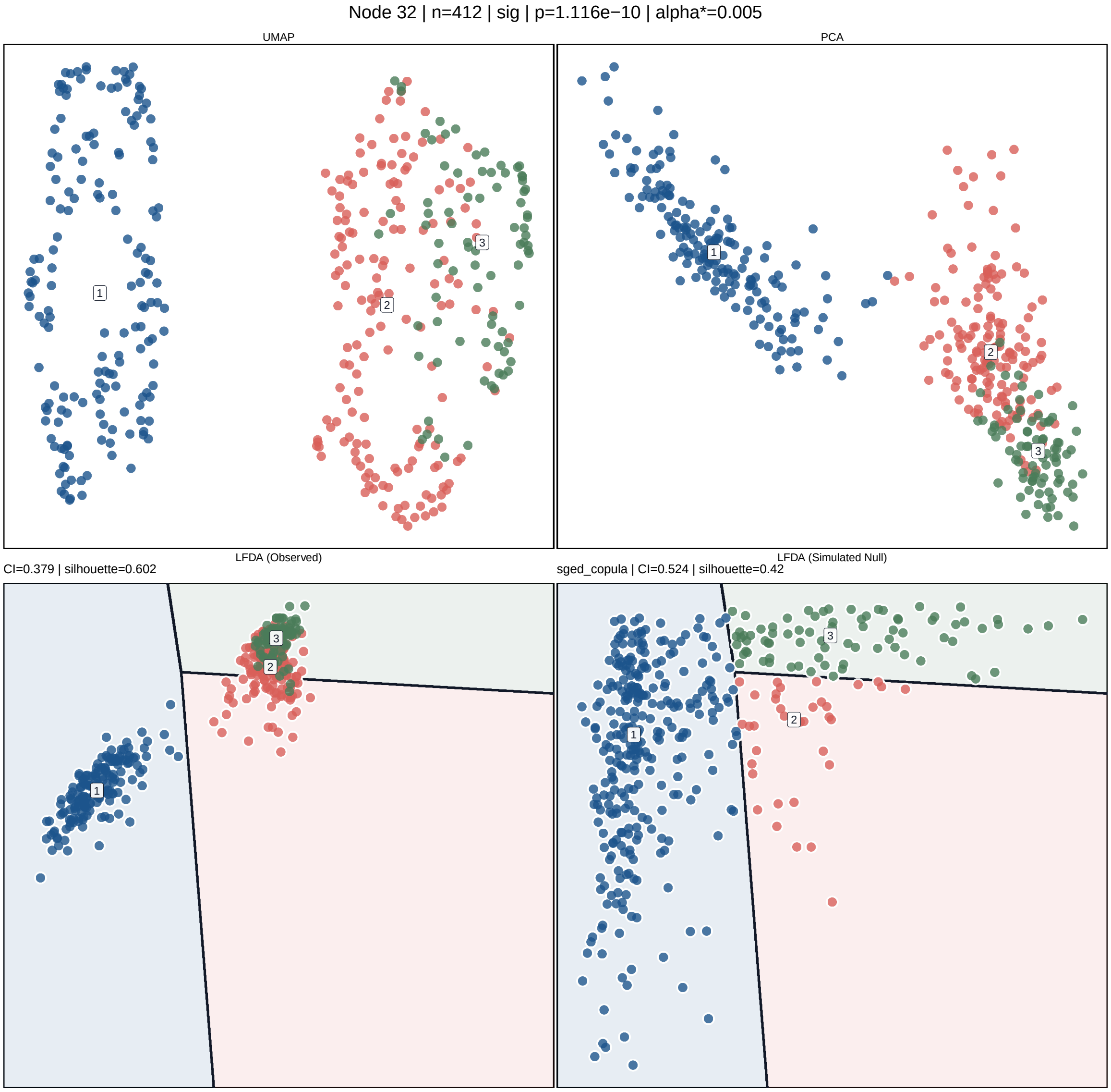


Figure S18: **Diagnostic visualization of node 32 in the Tabula Muris heart hierarchy.** Node 32 contained 412 cells from initial clusters C12, C13, and C15. Panels show the UMAP embedding (top left), PCA space (top right), observed two-dimensional LFDA projection (bottom left), and a simulated sample from the fitted null model (bottom right). Colors and numeric labels identify the three groups used for the observed-data visualization; simulated cells are colored by their assigned decision regions. Shaded backgrounds and black lines in the LFDA panels indicate nearest-centroid decision regions and their boundaries. The node-level test was significant ($p = 1.116 \times 10^{-10}$; $\alpha^* = 0.005$), and the hierarchy separated C12 from $\{C13, C15\}$. The displayed CI and silhouette values summarize the plotted samples.

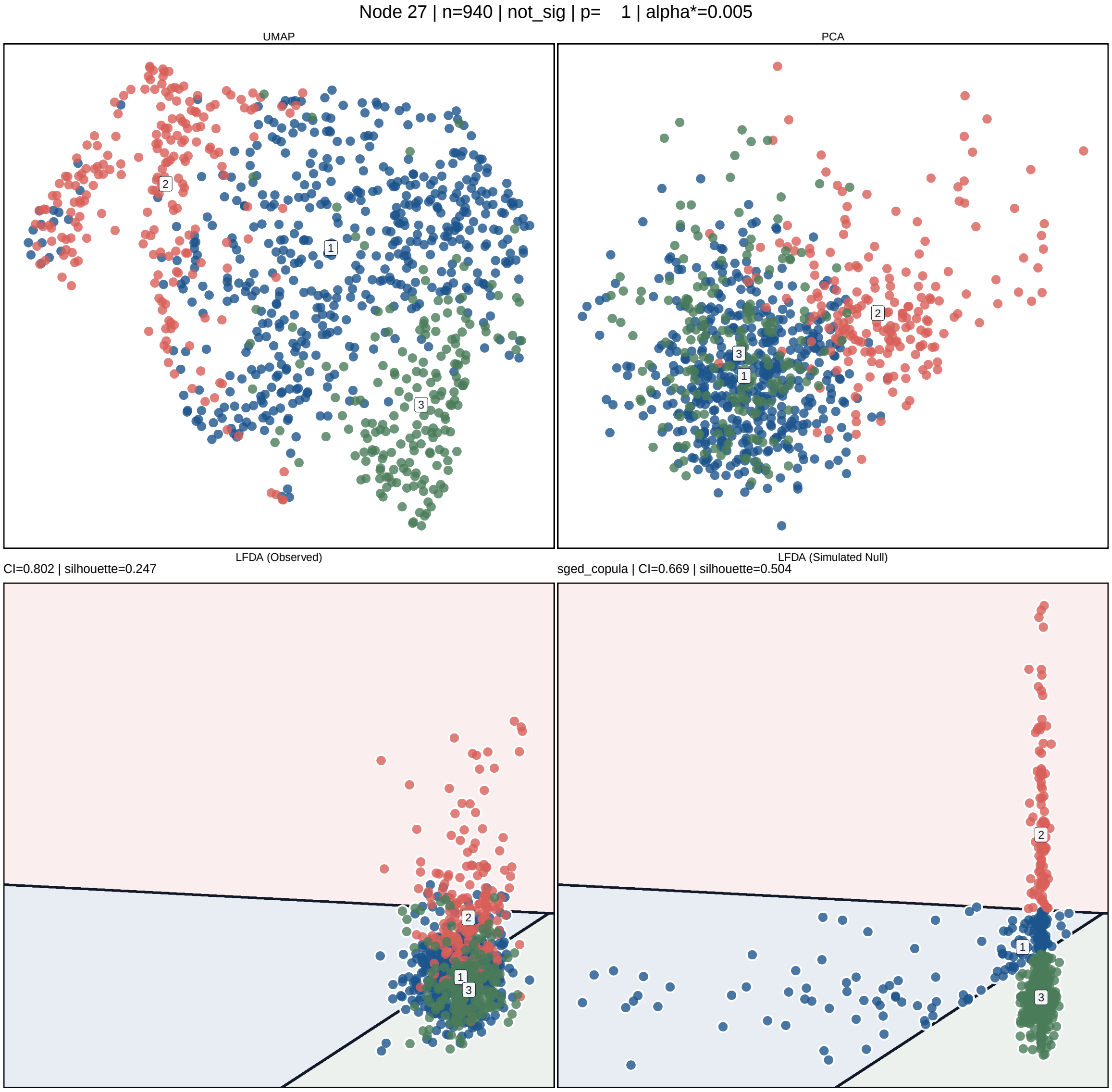


Figure S19: **Diagnostic visualization of node 27 in the Tabula Muris heart hierarchy.** Node 27 contained 940 cells from initial clusters C1, C7, and C9. Panels show the UMAP embedding (top left), PCA space (top right), observed two-dimensional LFDA projection (bottom left), and a simulated sample from the fitted null model (bottom right). Colors and numeric labels identify the three groups used for the observed-data visualization; simulated cells are colored by their assigned decision regions. Shaded backgrounds and black lines in the LFDA panels indicate nearest-centroid decision regions and their boundaries. The observed LFDA projection shows substantial overlap among the three groups. The node-level test was nonsignificant ($p = 1.000$; $\alpha^* = 0.005$), and C1, C7, and C9 were merged into a single final cluster. The displayed CI and silhouette values summarize the plotted samples.

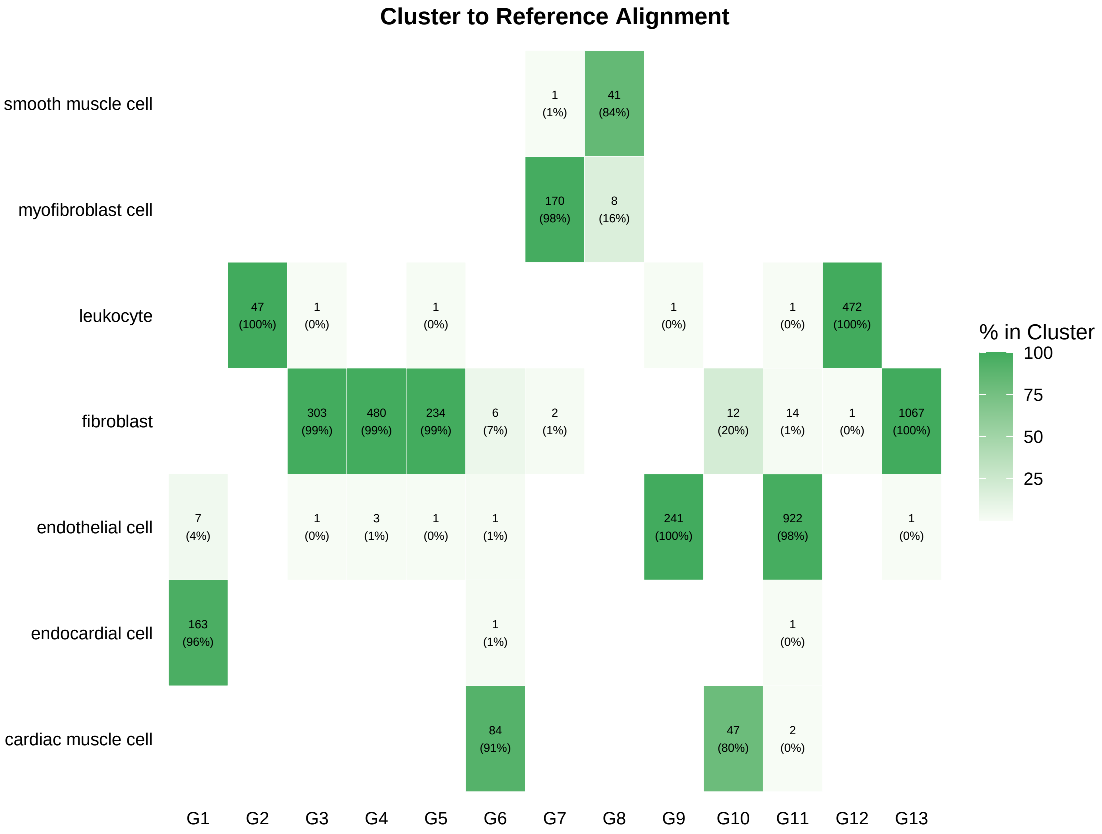


Figure S20: **Alignment between the final ProClust clusters and reference cell-type annotations in the Tabula Muris heart dataset.** Columns correspond to the 13 final ProClust clusters, and rows correspond to the reference cell-type annotations. Each nonempty cell reports the number of cells and the percentage of the corresponding ProClust cluster assigned to that reference annotation. Darker shading indicates a larger percentage within the ProClust cluster.

#### S5 Additional HLCA Results

**Additional clustering results.** We additionally ran Louvain and Leiden across resolutions 0.4–1.2 on the same HLCA immune dataset (Fig. S21); Leiden at resolution 1.0 is omitted here, as it corresponds to the over-clustered partition already shown in the main text. Monocle3 was run using its default number of nearest neighbors ($k = 20$) and default Leiden-based clustering algorithm. SC3 was not evaluated because its computational and memory requirements were high for the HLCA dataset.

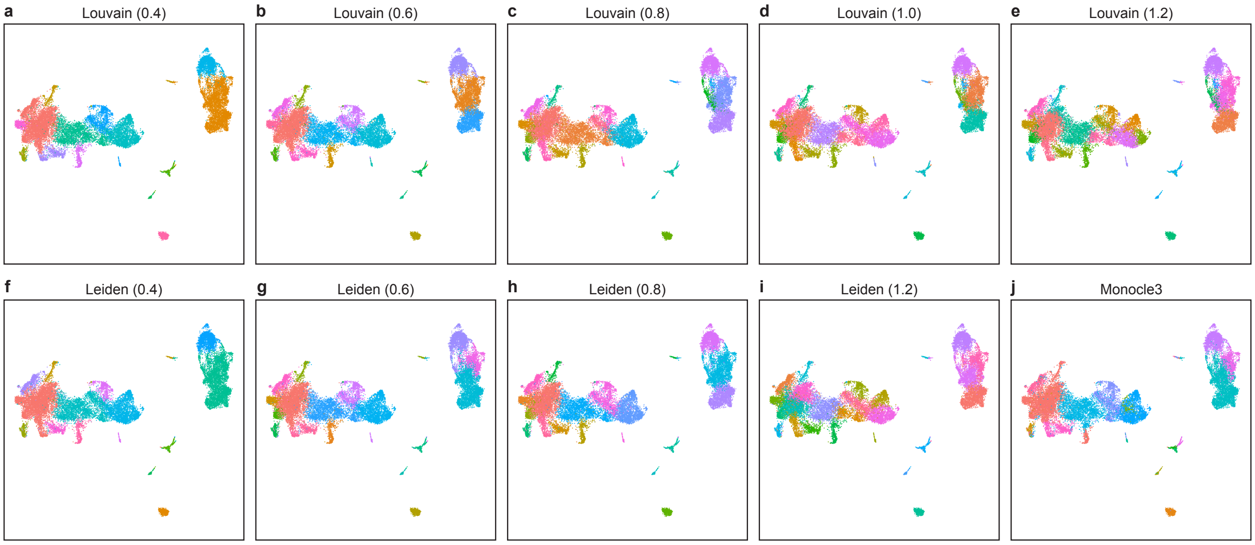


Figure S21: **Clustering results from Louvain, Leiden, and Monocle3 on the HLCA immune dataset. a–e**, UMAP embeddings colored by clusters identified by Louvain at resolutions 0.4, 0.6, 0.8, 1.0, and 1.2. **f–i**, UMAP embeddings colored by clusters identified by Leiden at resolutions 0.4, 0.6, 0.8, and 1.2. **j**, UMAP embedding colored by clusters identified by Monocle3.

**Alveolar macrophage subtype markers.** To examine heterogeneity within the alveolar macrophage population, we restricted to ProClust clusters whose dominant ground-truth annotation was alveolar macrophages and examined the expression of marker genes for previously described alveolar macrophage states (Fig. S22). For each state, up to five marker genes present in the dataset were selected from a predefined candidate list, and average expression per cluster was scaled across clusters. Cluster-specific patterns are discussed in the main text.

**CD4 and CD8 T-cell analysis.** We additionally considered a focused CD4/CD8 T-cell subset of the HLCA core dataset. Cells annotated as either CD4 T cells or CD8 T cells at `ann_level_4` and belonging to the Immune compartment at `ann_level_1` were retained, yielding 50,359 cells,

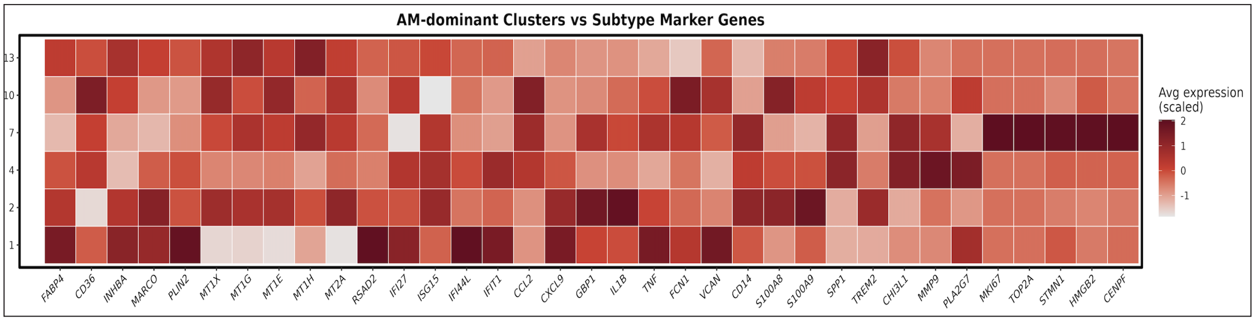


Figure S22: **Subtype-specific marker gene expression across alveolar macrophage clusters.** Heatmap showing the scaled average expression of marker genes associated with distinct alveolar macrophage states (homeostatic/lipid-associated, metallothionein, interferon-response, inflammatory, monocyte-associated, profibrotic, and proliferation-associated) across the subset of ProClust-identified clusters dominantly composed of alveolar macrophages.

including 21,285 CD4 T cells and 29,074 CD8 T cells. We specified a targeted panel of 44 T-cell marker genes representing CD4/CD8 lineage identity and major T-cell functional states; 42 of these genes were present in the expression matrix and retained for analysis.

Counts for the selected marker genes were normalized to a total expression of 10,000 per cell, log-transformed, and scaled. PCA was then performed using the 42 retained marker genes, yielding 41 principal components. Harmony was applied to the PCA representation using dataset identity as the batch variable. Potential outliers were identified in the Harmony-corrected representation using the Local Outlier Factor algorithm with 20 nearest neighbors and a contamination fraction of 0.05, removing 2,518 cells and leaving 47,841 cells.

A neighborhood graph was subsequently constructed from the Harmony-corrected representation using 30 nearest neighbors and cosine distance, followed by Leiden clustering at a resolution of 1.0, which produced 52 initial clusters for downstream hierarchical analysis.

The 52 Leiden clusters were used as the initial leaves for ProClust. A hierarchical tree was constructed by computing the centroid of each initial cluster in the Harmony-corrected representation and applying Ward's hierarchical clustering with the `ward.D2` linkage criterion to the Euclidean distances between cluster centroids. ProClust then proceeded recursively through this tree using the projected SigClust procedure. At each tested node, LFDA was used to obtain a node-specific discriminative representation, with at most three groups used in the LFDA projection. The skewed generalized error distribution copula (`sged_copula`) was used as the null model, and the cluster index was used as the test statistic. Testing used two-fold cross-validation, 20 clustering starts, and 200 Monte Carlo null replicates. The maximum number of principal components available for node-level dimension reduction was set to 10. Nodes without sufficient evidence of multimodality were merged, whereas significant nodes retained their descendant structure.

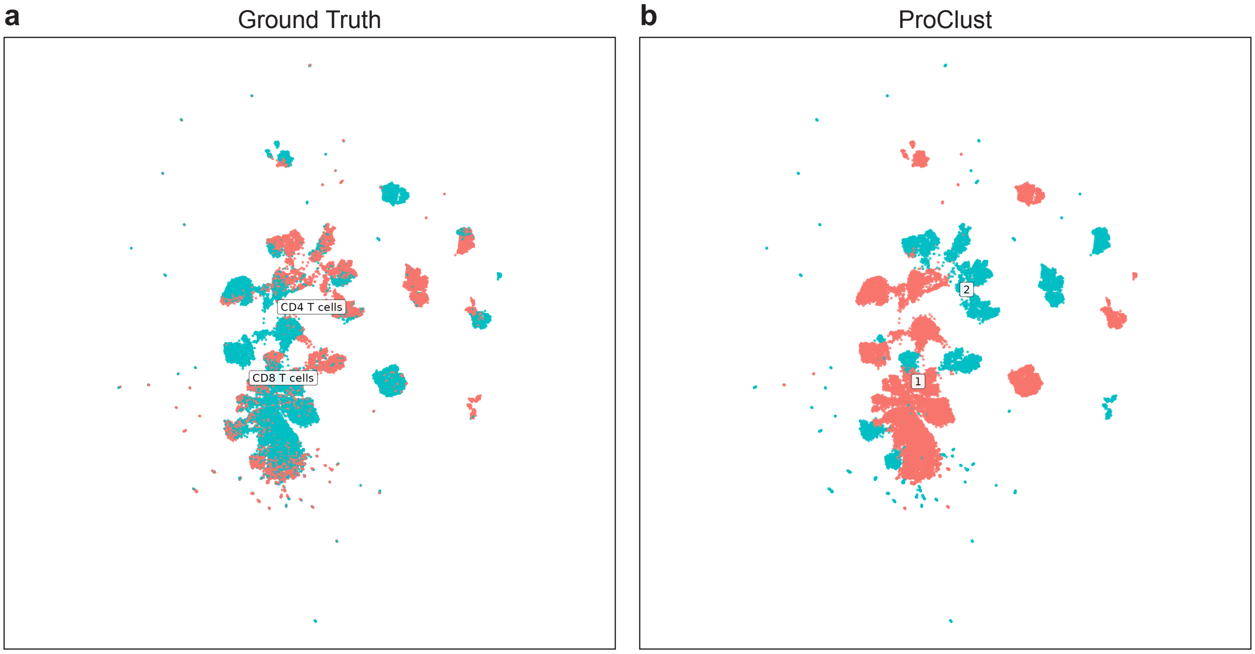


Figure S23: **Separation of CD4 and CD8 T cells using a targeted marker panel.** **a**, UMAP embedding of T cells colored by ground-truth CD4/CD8 annotations. **b**, UMAP embedding colored by clusters identified by ProClust using a targeted panel of T-cell lineage marker genes.

### S6 Additional Hydra Results

**Additional clustering results.** Across ectodermal and endodermal epithelial cells, ProClust recovered the major annotated cell states and showed comparable or higher agreement than most benchmark methods, as measured by AIR and NMI (Fig. S24).

**Neural/gland progenitor geometry depends on dimensionality reduction.** For neural/gland progenitor cells, substantial separation was apparent only in UMAP, whereas t-SNE, Harmony, and LFDA showed more continuous or overlapping distributions (Fig. S25). This dependence on the embedding suggests that the apparent gap may be introduced or exaggerated by UMAP rather than reflect a robust boundary in the underlying data.

**Marker expression in progenitor cells.** Hydra progenitor annotations were associated with the expression of selected marker genes: *HvSoxC* for SC/nb versus SC/prog, *Nowa-1* for nb3 versus

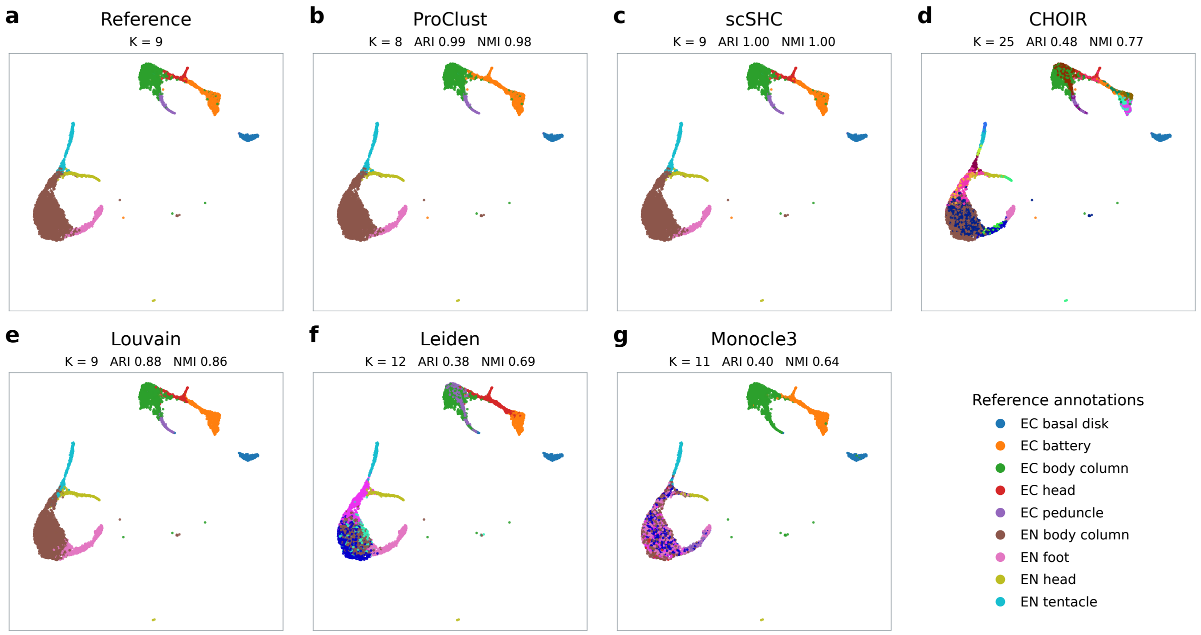


Figure S24: **Clustering of Hydra epithelial cells across methods.** UMAP representations of 6,742 epithelial cells – 2,520 ectodermal and 4,222 endodermal – colored by reference annotations (a), ProClust (b), scSHC (c), CHOIR (d), Louvain (e), Leiden (f), and Monocle3 (g).

nb5, and *ELAV2* for nc_gc_prog versus nc_prog (Fig. S26). Notably, overlapping expression and frequent non-detection indicate that these markers do not define unambiguous boundaries between the annotated states.

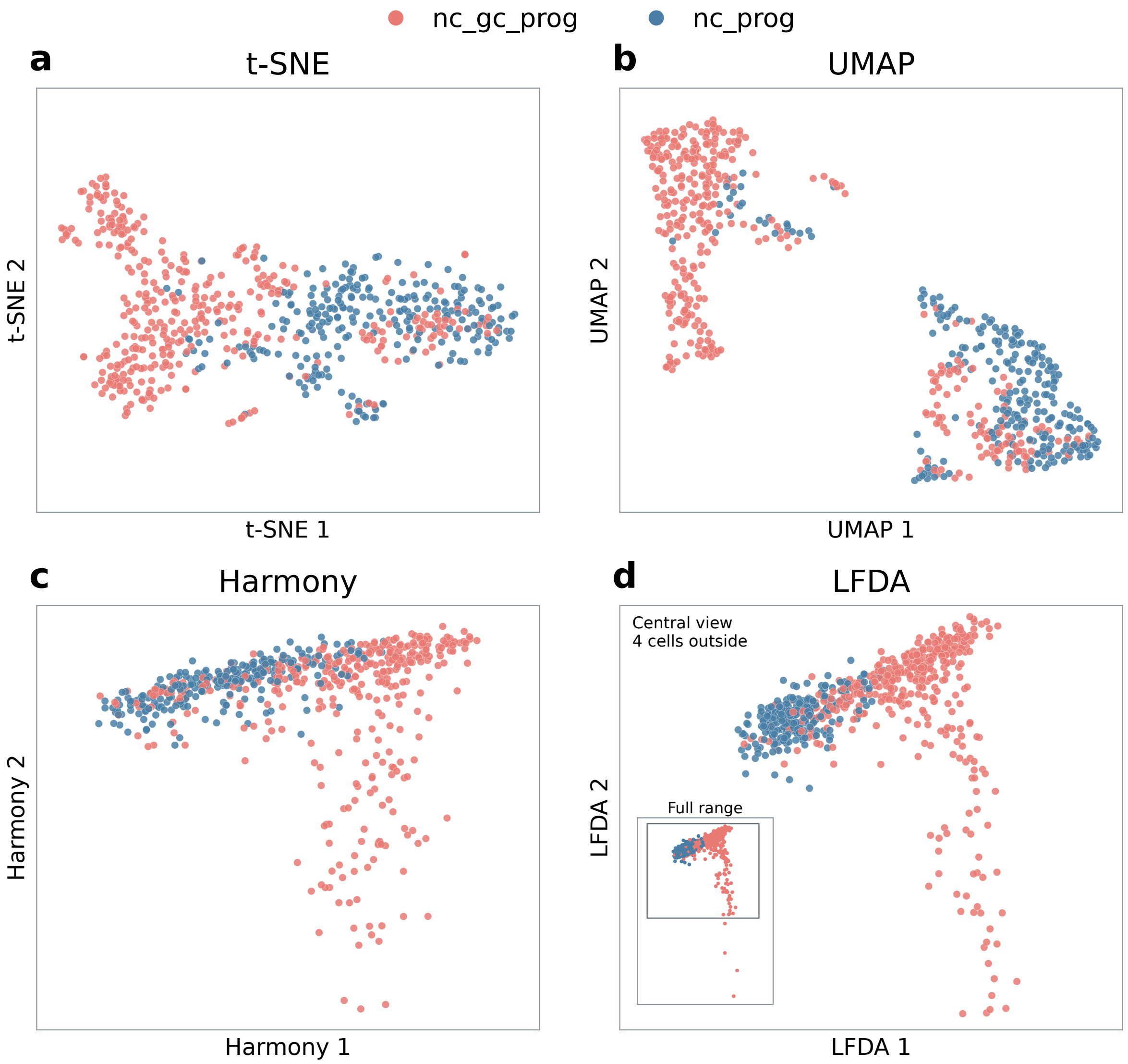


Figure S25: **Dependence of neural/gland progenitor geometry on dimensionality reduction.** The same 648 cells annotated as nc_gc_prog (n = 397) or nc_prog (n = 251) are displayed using t-SNE (a), UMAP (b), the first two Harmony coordinates (c), and LFDA (d).

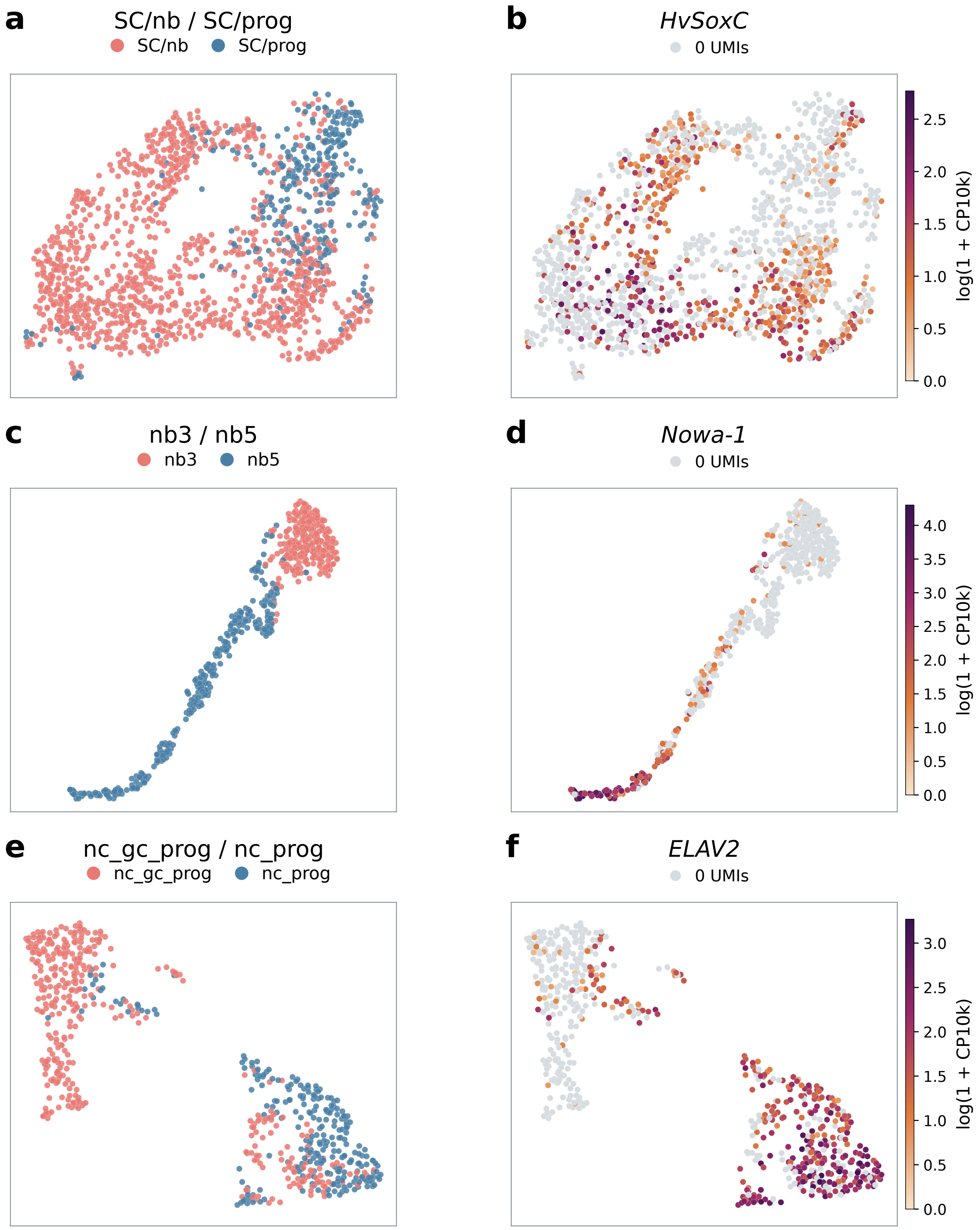


Figure S26: **Marker expression in relation to *Hydra* progenitor annotations.** UMAPs show reference annotations (a,c,e) alongside normalized expression (b,d,f) of HvSoxC in SC/nb and SC/prog cells (a,b; n = 1,546), Nowa-1 in nb3 and nb5 cells (c,d; n = 541), and ELAV2 in nc_gc_prog and nc_prog cells (e,f; n = 648). Expression is log normalized for depth. Gray indicates zero detected UMIs; positive-expression colors span each gene's full observed range.
